# The *Drosophila* chemokine-like Orion bridges phosphatidylserine and Draper in phagocytosis of neurons

**DOI:** 10.1101/2022.02.11.480148

**Authors:** Hui Ji, Bei Wang, Yifan Shen, David Labib, Joyce Lei, Xinchen Chen, Maria Sapar, Ana Boulanger, Jean-Maurice Dura, Chun Han

## Abstract

Phagocytic clearance of degenerating neurons is triggered by “eat-me” signals exposed on the neuronal surface. The conserved neuronal eat-me signal phosphatidylserine (PS) and the engulfment receptor Draper (Drpr) mediate phagocytosis of degenerating neurons in *Drosophila*. However, how PS is recognized by Drpr-expressing phagocytes *in vivo* remains poorly understood. Using multiple models of dendrite degeneration, we show that the *Drosophila* chemokine-like protein Orion can bind to PS and is responsible for detecting PS exposure on neurons; it is supplied cell-non-autonomously to coat PS-exposing dendrites and to mediate interactions between PS and Drpr, thus enabling phagocytosis. As a result, the accumulation of Orion on neurons and on phagocytes produces opposite outcomes by potentiating and suppressing phagocytosis, respectively. Moreover, the Orion dosage is a key determinant of the sensitivity of phagocytes to PS exposed on neurons. Lastly, mutagenesis analyses show that the sequence motifs shared between Orion and human immunomodulatory proteins are important for Orion function. Thus, our results uncover a missing link in PS-mediated phagocytosis in *Drosophila* and imply conserved mechanisms of phagocytosis of neurons.

**SIGNIFICANCE STATEMENT:** Phagocytes efficiently clear sick or damaged neuronal branches by engulfing them, while leaving healthy branches untouched. How phagocytes recognize degenerating neurites remains poorly understood. Here, we identified a key role for the secreted protein Orion in the detection and engulfment of degenerating neurites in *Drosophila*. Using multiple models of dendrite degeneration, we found that Orion acts as a bridging molecule between the neuronal “eat-me” signal phosphatidylserine and the engulfment receptor Draper on phagocytes, enabling phagocytosis. Our study reveals a missing link in phosphatidylserine-mediated phagocytosis *in vivo*, sheds light on factors determining the sensitivity of phagocytes, and implies the potential for manipulating the detection of neuronal “eat-me” signals in neurodegenerative diseases.

## INTRODUCTION

Phagocytosis of apoptotic and degenerative neurons is essential for the development and homeostasis of the nervous system (1, 2). Abnormal phagocytosis is also associated with neuroinflammation and neurodegenerative diseases (3). Neuronal debris is recognized and cleared by resident phagocytes of the nervous system through “eat-me” signals exposed on the neuronal surface. A conserved “eat-me” signal is phosphatidylserine (PS), a negatively charged phospholipid normally kept in the inner leaflet of the plasma membrane by P4-ATPase flippases (4). During neurite degeneration and apoptosis, PS is externalized to the outer surface of neuronal membranes (5–8). Exposed PS dominantly triggers phagocytosis of neurons, as demonstrated by the observation that loss of PS flippases in neurons results in PS exposure and neurodegeneration across species (6, 9, 10). Besides clearing existing neuronal debris, PS-mediated phagocytosis also drives the degeneration of injured neurites and neurons with certain genetic perturbations (6, 11). In the central nervous system (CNS), local PS exposure enables microglia-mediated synaptic elimination (12–14). Thus, the regulation and recognition of neuronal PS exposure are critical for the development and homeostasis of the nervous system.

*Drosophila* has been an important model organism for studying neuronal phagocytosis. In *Drosophila,* Draper (Drpr) is the best-known receptor responsible for phagocytosis of neurons (15). As a homolog of the *C. elegans* engulfment receptor CED-1 (16) and the mammalian engulfment receptors Jedi-1 and MEGF10 (17), Drpr is involved in many contexts of neuronal phagocytosis, including the clearance of apoptotic neurons during embryonic development (15, 18), axon and dendrite pruning during neuronal remodeling (19, 20), injury-induced neurite degeneration (21, 22), and removal of destabilized boutons at neuromuscular junctions (23). Despite the well-known importance of Drpr in sculping the nervous system, how Drpr recognizes degenerating neurons *in vivo* is still unclear.

Recently, the secreted protein Orion was discovered as being required for the developmental pruning and clearance of *Drosophila* mushroom body (MB) axons (24). Orion shares a CX_3_C motif with mammalian CX3CL1 (also known as fractalkine), which is required for the elimination of synapses in the mouse barrel cortex (25). CX3CL1 is known as a chemokine because of its ability to direct migration of leukocytes and microglia (26, 27). Thus, Orion represents the first known chemokine-like molecule in *Drosophila* and shares conserved functions with CX3CL1 in the remodeling of the nervous system. However, how Orion and CX3CL1 exactly function in the phagocytosis of neurons is still unknown.

In this study, we examined Orion’s function in the phagocytosis of *Drosophila* class IV dendritic arborization (C4da) neurons, a well-established *in vivo* model for studying PS-mediated phagocytosis (6, 11, 28). Here we present *in vivo* evidence that the transmembrane engulfment receptor Drpr relies on Orion to sense PS on neurons. Orion is secreted by peripheral non-neural tissues and serves as a non-autonomous permissive factor for phagocytosis of neurons. Orion binds to PS-exposing neurons and is also enriched on phagocytes over-expressing Drpr. Strikingly, a membrane-tethered Orion expressed by neurons dominantly induces phagocytosis even in the absence of PS exposure, while Orion accumulation on the surface of phagocytes makes phagocytes blind to PS-exposing neurons. Importantly, the dosage of Orion determines the sensitivity of phagocytes to neuronal PS exposure. Lastly, we establish that the motifs Orion shares with human chemokines and neutrophil peptides are critical for engulfment of PS-exposing neurons. These findings reveal key mechanisms of PS recognition in *Drosophila* and imply potentially conserved roles of chemokines in PS-mediated phagocytosis of neurons.

## RESULTS

### *orion* is required for phagocytosis of dendrites

To determine if *orion* is involved in phagocytosis of degenerating dendrites of da neurons, we first examined phagocytosis of injured C4da dendrites in the *orion^1^* mutant, which results in a G to D mutation in the C-termini of both Orion protein isoforms (Figure S1A) and abolishes clearance of pruned axons during MB remodeling (24). C4da neurons grow elaborate sensory dendrites on larval epidermal cells (Han et al., 2012), which act as the primary phagocytes during dendrite degeneration and remodeling (Han et al., 2014). Dendrites were severed from the cell body using laser, and engulfment of the injured dendrites was visualized using MApHS, a pH-sensitive dendritic marker consisting of extracellular pHluorin and intracellular tdTom (28). Engulfment was signified by the loss of pHluorin signal due to the drop of pH in early phagosomes (28, 29). Injured dendrites of C4da neurons in the control larvae were completely engulfed by epidermal cells 24 hours (hrs) after injury (AI), as indicated by the loss of pHluorin signals on tdTom-positive dendritic debris dispersed in epidermal cells (Figures 1A-1A”, and 1D). Because *orion* is located on the X chromosome, we examined both hemizygous male larvae and homozygous female larvae of *orion^1^*. In contrast to the wildtype (WT), both groups of *orion^1^* mutants showed little engulfment of injured dendrites by epidermal cells (Figures 1B-1C”, and 1D), as indicated by the presence of pHluorin signals on tdTom-labeled debris of injured dendrites. The debris remained in the original dendritic patterns, which is another sign of impaired engulfment by epidermal cells. These results suggest that *orion* is required for the engulfment of injured dendrites. Since male and female *orion* mutants exhibited similar degrees of engulfment defects (Figure 1D), we used hemizygous male mutants only in subsequent assays.

**Figure 1:**
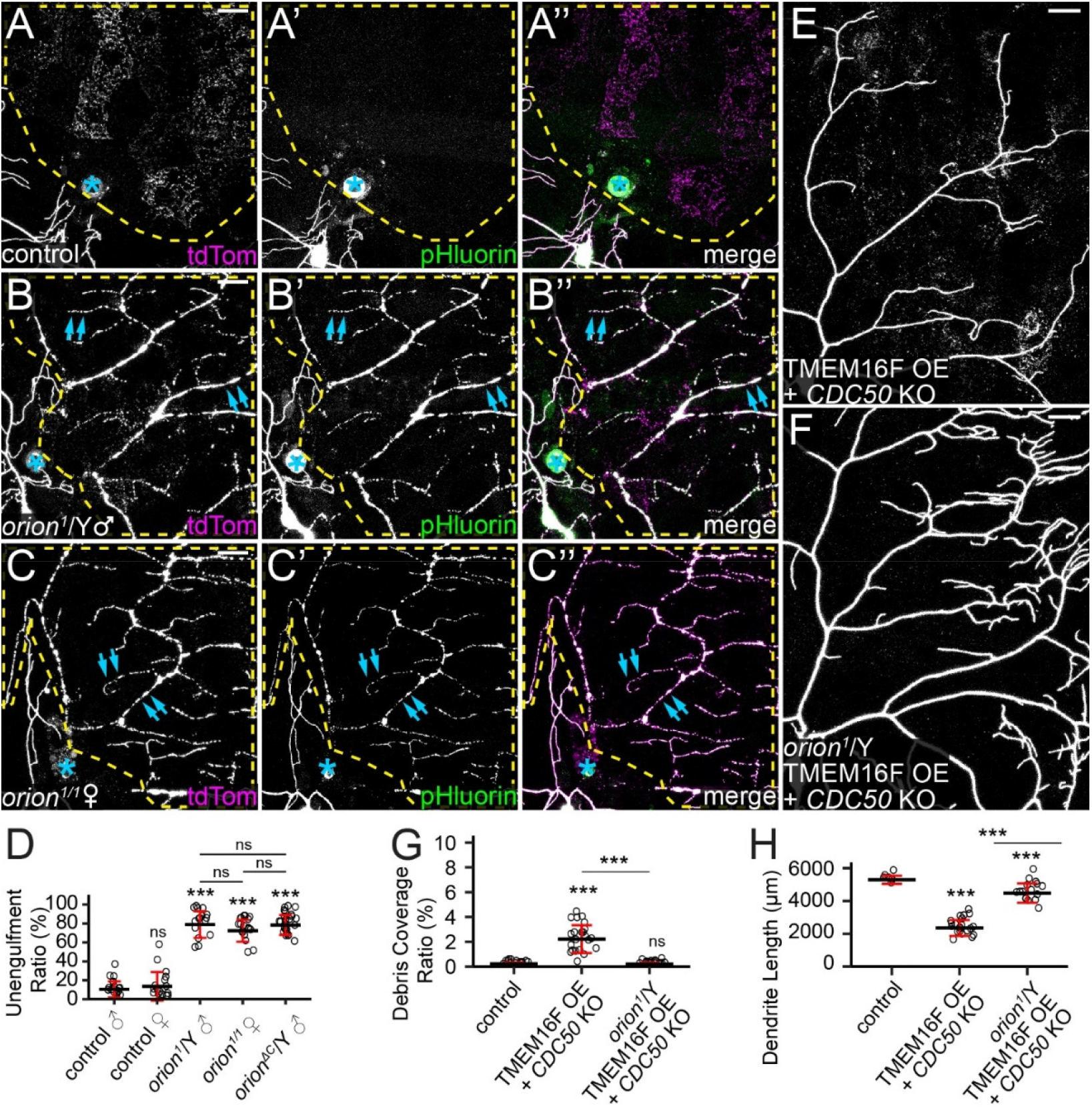
*orion* is required for phagocytosis of dendrites in multiple dendrite degeneration paradigms. (A-C”) Partial dendritic fields of ddaC neurons in control (A-A”), *orion^1^* hemizygous (B-B”), and *orion^1^* homozygous (C-C”) larvae at 22-24 hrs after injury (AI). Yellow dash outlines: territories originally covered by injured dendrites; blue asterisks: injury sites; blue arrows: injured but unengulfed dendrite fragments. (D) Quantification of unengulfment ratio of injured dendrites (pHluorin-positive debris area/tdTom-positive debris area). n = number of neurons and N = number of animals: control males (n = 18, N = 10); control females (n = 16, N = 9); *orion^1^*/Y (n = 16, N = 8); *orion^1/1^* (n = 18, N = 9); *orion^ΔC^*/Y (n = 25, N = 12). One-way ANOVA and Tukey’s test. (E and F) Partial dendritic fields of TMEM16F OE +*CDC50* KO neurons in the wildtype background (E) and *orion^1^* hemizygous background (F). (G and H) Quantification of debris coverage ratio (percentage of debris area normalized by dendrite area) (G) and dendrite length (H) at 140 hrs after egg laying (AEL). n = number of neurons and N = number of animals: control (n = 14, N = 7); TMEM16F OE +*CDC50* KO (n = 22, N = 11); TMEM16F OE +*CDC50* KO in *orion^1^*/Y (n = 16, N = 8). For (G), Kruskal-Wallis (One-way ANOVA on ranks) and Dunn’s test, p-values adjusted with the Benjamini-Hochberg method; for (H), One-way ANOVA and Tukey’s test. In all image panels, neurons were labeled by *ppk-MApHS*. Scale bars, 25 μm. For all quantifications, ***p≤0.001; n.s., not significant. The significance level above each genotype is for comparison with the control. Black bar, mean; red bar, SD. See also Figure S1.

In addition to phagocytosis after injury, C4da dendrites are all pruned and engulfed by epidermal cells during metamorphosis (28). Consistent with the role of *orion* in axonal pruning of MB neurons (24), we also confirmed that *orion^1^* exhibits strong defects in the clearance of pruned dendrites of C4da neurons during metamorphosis (Figure S1B-S1D).

The engulfment of injured dendrites by epidermal cells is mediated by PS exposure on dendrites (6, 11). To test directly if *orion* is required for PS exposure-induced phagocytosis, we examined a paradigm in which PS exposure was ectopically induced in otherwise healthy neurons using a combination of *CDC50* knockout (KO) and TMEM16F overexpression (OE). *CDC50* encodes a chaperone protein required for the activity of P4-ATPases that keep PS in the inner leaflet of the plasma membrane (30). TMEM16F is a mammalian scramblase that mixes PS between the two leaflets of the plasma membrane (31, 32). Dendrites with *CDC50* KO and TMEM16F OE expose PS and shed membrane in a phagocytosis-dependent manner (6). Whereas these neurons showed reduced dendrite length and elevated debris levels in wandering 3^rd^ instar larvae with wildtype *orion* (Figures 1E, 1G, and 1H), *orion^1^* hemizygous males exhibited almost normal dendritic length and no dendritic debris in epidermal cells (Figures 1F-1H), indicating a lack of engulfment by phagocytes.

Because *orion^1^* carries a missense mutation that may not completely abolish Orion function, we compared its properties to those of *orion^ΔC^*, a predicted null mutation that lacks all three common C-terminal exons of both *orion* isoforms (Figure S1A) (24). *orion^ΔC^* showed similar levels of phagocytosis defects as *orion^1^* in injury-induced degeneration (Figures S1E-S1E”, and 1D), suggesting that *orion^1^* has lost most, if not all, of its function. Since *orion* encodes two isoforms with different N-terminal sequences (Figure S1A), we next asked if one or both isoforms contribute to phagocytosis. Mutations targeting each of the two *orion* isoforms separately (Figure S1A) showed no phagocytosis defects in injury-induced degeneration (Figure S1F-S1G”), suggesting that OrionA and OrionB isoforms are redundant in the phagocytosis of injured dendrites. These results are consistent with the redundant effects of the same mutations in MB neuron axon pruning (24).

### Orion functions cell-non-autonomously

*orion* encodes a secreted protein, raising the question of how it contributes to phagocytosis of dendrites. To determine which tissues express Orion, we generated an *orion* knock-in (KI) allele in which the C-terminus of *orion* is fused in-frame with 4 copies of mNeonGreen2_11_ (mNG2_11_) (33), a V5 tag, a self-cleaving F2A sequence (34), and LexA::VP16 (35) (Figure S2A). This allele, abbreviated as *orion^KI^*, has several purposes: detection of *orion*-expressing cells by LexA::VP16 transcription activity, detection of endogenous Orion proteins by V5 staining, and visualization of endogenous Orion proteins in a tissue-specific manner by reconstitution of mNG2 fluorescence with tissue-specific expression of mNG2_1-10_ (33).

By using *orion^KI^* with a *LexAop-GFPnls* or a *LexAop-GFP* reporter, we found that *orion* transcripts are expressed in many peripheral tissues in larvae, including epidermal cells, hemocytes, trachea, and the fat body (Figure 2A-2B). However, *orion* transcription is missing in larval da neurons (Figure 2A, inset), even though a weak *orion* transcription activity was later detected in a subset of da neurons at 5-12 hrs after puparium formation (APF) (Figures S2B-S2B”), when some da neurons die and others undergo dendritic pruning (36). In the central nervous system, we found that *orion* was transcribed in most glial cells and in a small subset of neurons at the wandering 3^rd^ instar larval stage (Figure S2C-S2D”). Consistent with a role of *orion* in axonal pruning of MB neurons, *orion* is expressed in Kenyon cells at 8 hrs APF (Figure S2E). Despite the robust LexA::VP16 activity in *orion^KI^*, Orion proteins appear to be expressed at a low level. Using V5 staining, we could observe only weak Orion signals on the larval epidermis (Figures S2F and S2G). By expressing secreted mNG2_1-10_ in a patch of epidermal cells (in the R16D01 domain), we also detected weak Orion-mNG2 signals in epidermal cells (Figures S2H and S2I).

**Figure 2:**
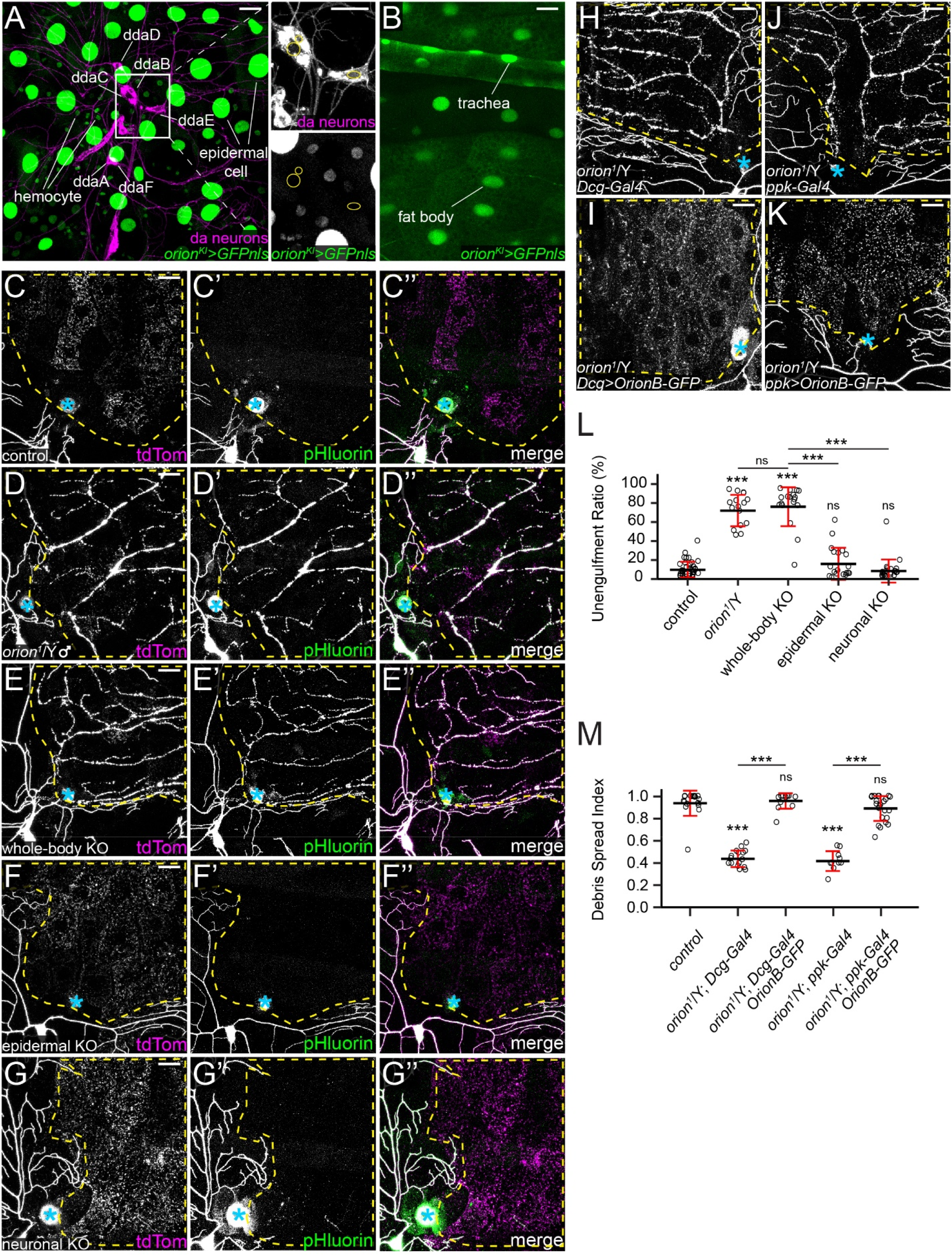
Orion functions cell-non-autonomously. (A-B) GFPnls expression driven by *orion-LexA* (*orion^KI^*) in epidermal cells, hemocytes (A), trachea, and fat body (B), but not in da neurons (A, insets). Yellow circles indicate nuclei of ddaB, ddaC, and ddaE neurons (A). Scale bars, 25 μm (A), 10 μm (A insets), and 50 μm (B). (C-G”) Partial dendritic fields of neurons in control (C-C”, same images as Figure 1A-1A”), *orion^1^* hemizygotes (D-D”, same images as Figure 1B-1B” for clarity), whole-body *orion* KO (E-E”), epidermal *orion* KO (F-F”), and neuronal *orion* KO (G-G”) at 22-24 hrs AI. (H-K) Partial dendritic fields of ddaC neurons in *orion^1^* hemizygotes with fat body Gal4 (H), with OrionB-GFP OE in the fat body (I), with neuron Gal4 (J), and with OrionB-GFP OE in neurons (K) at 22-24 hrs AI. (L) Quantification of unengulfment ratio of injured dendrites. n = number of neurons and N = number of animals: control and *orion^1^*/Y (same dataset as in Figure 1D); whole-body *orion* KO (n = 18, N = 10); epidermal *orion* KO (n = 19, N = 11); neuronal *orion* KO (n = 21, N = 11). One-way ANOVA and Tukey’s test. (M) Quantification of debris spread index of injured dendrites (area ratio of all 15 pixel x 15 pixel squares that contained dendrite debris in the region of interest). n = number of neurons and N = number of animals: *orion^1^*/Y (n = 18, N =10); *Dcg-Gal4* in *orion^1^*/Y (n = 16, N = 9); *Dcg>OrionB-GFP* in *orion^1^*/Y (n = 13, N = 7), *ppk-Gal4* in *orion^1^*/Y (n = 10, N = 6); *ppk>OrionB-GFP* in *orion^1^*/Y (n = 24, N = 12). One-way ANOVA and Tukey’s test. Neurons were labeled by *21-7*>*UAS-CD4-tdTom* (A-B),*ppk-MApHS* (C-G”), *ppk-CD4-tdTom* (H, I), and *ppk*>*CD4-tdTom* (J, K). In (C-K), yellow dash outlines: territories originally covered by injured dendrites; blue asterisks: injury sites; scale bars, 25 μm. For all quantifications, ***p≤0.001; n.s., not significant. The significance level above each genotype is for comparison with the control. Black bar, mean; red bar, SD. See also Figure S2.

To investigate whether *orion* is required in specific tissues for phagocytosis of injured dendrites, we knocked out *orion* in either neurons or epidermal cells using tissue-specific Cas9s and ubiquitously expressed guide RNAs (gRNAs) (37). As a control, knocking out *orion* with a ubiquitous Cas9 (*Act5C-Cas9*) (38) faithfully replicated the phagocytosis defects of *orion^1^* (Figures 2C-2E”, and 2L), demonstrating the effectiveness of *orion* KO. However, *orion* KO in either da neurons alone (with *SOP-Cas9*; (37)) or in epidermal cells alone (with *shot-Cas9*; (11)) did not interfere with the engulfment of injured dendrites (Figures 2F-2G”, and 2L).

The lack of effect of *orion* KO in da neurons or in epidermal cells suggests that Orion functions cell-non-autonomously. We further tested this idea by asking whether supplying Orion in the extracellular space is sufficient to rescue the impaired engulfment of injured dendrites in *orion* mutants. Extracellular Orion supply was achieved by overexpressing Orion in the fat body, which can efficiently secrete proteins into the hemolymph (6). Because overexpression (OE) of OrionA in the fat body caused early larval lethality, we chose to overexpress a GFP-tagged OrionB (OrionB-GFP) in these rescue experiments. While the debris of injured dendrites remained in the original dendrite patterns in *orion^1^* hemizygotes (Figure 2H), which indicates a lack of engulfment by epidermal cells, OrionB overexpression in the fat body of *orion^1^* hemizygotes resulted in even spreading of the debris in the underlying epidermal cells (Figure 2I), suggesting successful rescue of engulfment. Similarly, neuronal expression of OrionB-GFP also fully rescued the engulfment defects as measured by the spread of debris (Figure 2J, 2K, and 2M).

Together, these results indicate that Orion is primarily expressed in non-neural tissues in the periphery and functions cell-non-autonomously for the engulfment of dendrites.

### Phosphatidylserine exposure induces Orion binding to the cell surface

To further understand the engulfment defects in *orion* mutants, we examined whether PS exposure on injured dendrites is affected in *orion* mutants. We used fat body-derived Annexin V-mCardinal (AV-mCard) as a PS sensor; it labels injured but not healthy dendrites (6). AV-mCard robustly labeled dendrite fragments in *orion^ΔC^* mutant larvae at 24 hrs AI (Figure 3A-3B), suggesting that *orion* LOF does not interfere with PS exposure on injured dendrites.

**Figure 3:**
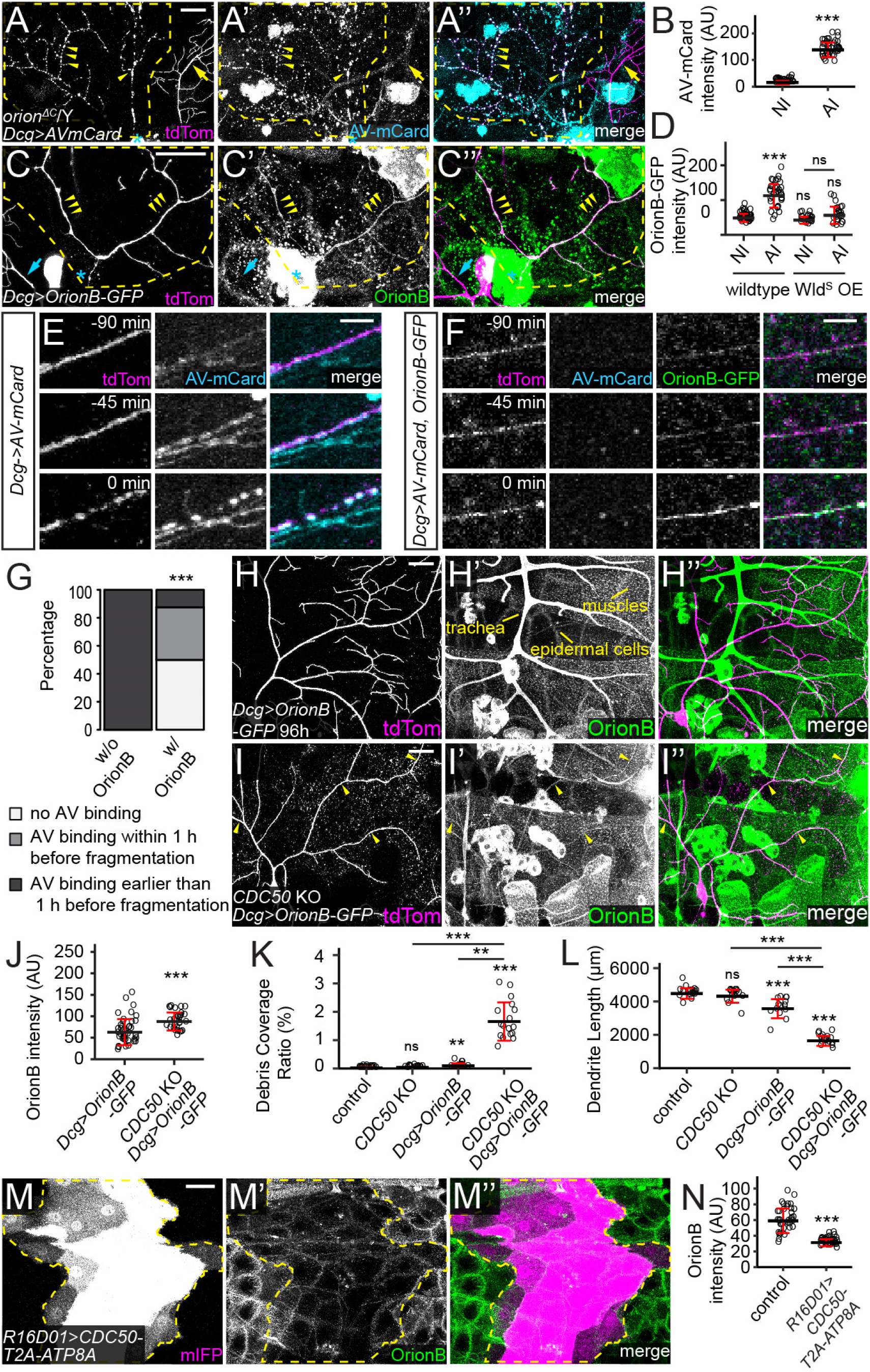
PS exposure induces Orion binding to the cell surface. (A-A”) Labeling of injured dendrites of a ddaC neuron by AV-mCard at 23 hrs AI in *orion^ΔC^* hemizygous larvae. Yellow arrowheads: injured dendrites with AV-mCard labeling; yellow arrows: uninjured dendrites lacking AV-mCard labeling. (B) Quantification of AV-mCard binding on dendrites. AV-mCard intensities were measured on both uninjured (NI) and injured (AI) dendrites. n = number of measurements and N = number of animals: NI (n = 39, N = 8); AI (n = 39, N = 8). Welch’s t-test. (C-C”) Labeling of injured dendrites of a ddaC neuron by Orion-GFP at 6 hrs AI in wildtype larvae. Yellow arrowheads: injured dendrites with OrionB-GFP labeling. (D) Quantification of Orion-GFP binding on dendrites of wildtype and Wld^S^ OE neurons. n = number of measurements and N = number of animals: wildtype NI (n = 39, N = 10); wildtype AI (n = 36, N = 10); Wld^S^ OE NI (n = 22, N = 6); Wld^S^ OE AI (n = 23, N = 6). Kruskal-Wallis (One-way ANOVA on ranks) and Dunn’s test, p-values adjusted with the Benjamini-Hochberg method. (E-F) Time series of injured dendrites of ddaC neurons from 90 min before fragmentation to the moment of fragmentation, with only AV-mCard expressed (E) or both OrionB-GFP and AV-mCard co-expressed (F) by the fat body. Time stamps are relative to the frame of dendrite fragmentation. (G) Percentages of injured dendrites showing no AV binding, AV binding within 1 hr before fragmentation, AV binding earlier than 1 hr before fragmentation. n = number of measurements and N = number of animals: w/o OrionB OE (n = 9, N = 5); w/ OrionB OE (n = 8, N = 3). Fisher’s exact test. (H-I”) Distribution of fat body-derived OrionB-GFP with wildtype (H-H”) and *CDC50* KO (I-I”) dendrites at 96 hrs AEL. Peripheral tissues showing OrionB-binding are labeled in (H’). Yellow arrowheads indicate OrionB-binding on *CDC50* KO dendrites (I-I”). (J) Quantification of OrionB-GFP binding on wildtype and *CDC50* KO dendrites. n = number of measurements: *Dcg*>*OrionB-GFP* (n = 41, N = 7); *CDC50* KO +*Dcg*>*OrionB-GFP* (n = 29, N = 6). Welch’s t-test. (K-L) Quantification of debris coverage ratio (K) and dendrite length (L) at 96 hrs AEL. n = number of neurons and N = number of animals: control (n = 17, N = 9); *CDC50* KO (n = 15, N = 8); *Dcg*>*OrionB-GFP* (n = 13, N = 8); *CDC50* KO + *Dcg*>*OrionB-GFP* (n = 17, N = 9). For (K), Kruskal-Wallis (One-way ANOVA on ranks) and Dunn’s test, p-values adjusted with the Benjamini-Hochberg method; for (L), one-way ANOVA and Tukey’s test. (M-M”) OrionB-GFP binding on epidermal cells that expressed *CDC50-T2A-ATP8A*. Yellow dash outlines: *CDC50-T2A-ATP8A* overexpressing region. (N) Quantification of OrionB-GFP binding on wildtype epidermal cells and *CDC50-T2A-ATP8A* OE epidermal cells. n = number of measurements and N = number of animals: control (n = 36, N = 9); *R16D01*>*CDC50-T2A-ATP8A* (n = 36, N = 9). Welch’s t-test. In (A-A”) and (C-C”), yellow dash outlines: territories originally covered by injured dendrites; blue asterisks: injury sites. Neurons were labeled by *ppk-MApHS* (A-A” and E-E”), and *ppk-CD4-tdTom* (C-C”, F-F” and H-I”). Scale bars, 25 μm (A-A”, C-C”, H-I”), 5 μm (E-F), and 50 μm (M-M”). For all quantifications, **p≤0.01, ***p≤0.001; n.s., not significant. The significance level above each genotype is for comparison with the control. Black bar, mean; red bar, SD. See also Movie S1-S3 and Figure S3. **Movie S1**: OrionB-GFP labels degenerating dendrites after injury, related to Figure 3. Time-lapse movie of laser-injured ddaC dendrites from 2.5 to 11 hrs AI, showing OrionB-GFP labeling on injured dendrites. The dendrite arbor on the bottom right was injured while the ones at the top were not. The mobile cells in the OrionB-GFP channel are hemocytes. Time stamp is relative to the first frame. **Movie S2**: AV-mCard labels injured dendrites before dendrite fragmentation, related to Figure 3. Time-lapse movie of laser-injured ddaC dendrites from 2.5 to 5 hrs AI, showing AV-mCard labeling on injured dendrites as early as 2 hrs before fragmentation. The AV-mCard signals that do not colocalize with tdTom signals indicate AV-mCard labeling on injured dendrites of other types of neurons. Time stamp is relative to the frame of dendrite fragmentation. **Movie S3**: AV-mCard fails to label injured dendrites when OrionB-GFP is co-expressed, related to Figure 3. Time-lapse movie of laser-injured ddaC dendrites from 1 to 3 hrs AI showing OrionB-GFP labeling but not AV-mCard labeling on injured dendrites before fragmentation. Time stamp is relative to the frame of dendrite fragmentation.

Because Orion is a secreted protein, we asked where Orion is located during engulfment. We observed strong enrichment of fat body-derived OrionB-GFP on injured dendrites at 4-6 hrs AI (Figure 3C-3D, and Movie S1), a timepoint when injured dendrites expose high levels of PS (6). Overexpression of Wld^S^, a transgene that suppresses fragmentation and PS exposure of injured dendrites when overexpressed in neurons (6, 11), efficiently suppressed OrionB-GFP enrichment on injured dendrites (Figures S3A-S3A”, and 3D), indicating that PS exposure may be required for Orion binding to injured dendrites.

To test the affinity of Orion for PS-exposing injured dendrites *in vivo*, we examined whether OrionB can compete with the PS sensor AV-mCard for binding to injured dendrites, given that AV binds to PS directly (4). Using long-term time-lapse imaging (39), we found that fat body-derived AV-mCard accumulated on injured dendrites at least 60 minutes before dendrite fragmentation (9/9 movies) (Figures 3E and 3G, and Movie S2). However, when both OrionB-GFP and AV-mCard were co-expressed by the fat body, OrionB-GFP was detected on injured dendrites long before fragmentation (8/8 movies), while AV-mCard did not bind the same injured dendrite in half of the cases (4/8 movies) (Figures 3F and 3G, and Movie S3). In most cases where AV-mCard was observed on injured dendrites (3/4 movies), the labeling only appeared right at the time of dendrite fragmentation (Figure 3G), when PS exposure is at its peak (6). These results suggest that Orion has a higher affinity to injured dendrites compared with AV.

To test if neuronal PS exposure is sufficient to induce Orion binding, we ectopically induced dendritic PS exposure by knocking out *CDC50* in neurons (6). Distribution of fat body-derived OrionB-GFP was examined at 96 hrs AEL, a time when *CDC50* KO alone is not yet sufficient to cause membrane loss of dendrites (Figures 3K and 3L). OrionB-GFP showed little binding to wildtype dendrites (Figures 3H-3H”, and 3J) but bound robustly to *CDC50* KO dendrites (Figures 3I-3J) at this stage. Interestingly, the presence of OrionB-GFP caused appreciable degeneration of *CDC50* KO dendrites, as indicated by the drastically increased dendrite debris and shortened total dendrite length (Figures 3K and 3L). These results suggest that Orion is recruited to PS-exposing dendrites and OrionB binding on dendrites potentiates epidermal engulfment.

The PS-binding C1C2 domain of mouse Lactadherin (LactC1C2) has been used as a PS sensor in multiple systems (5, 6, 40, 41). We previously reported that fat body-derived LactC1C2 not only labels degenerating dendrites but also promotes degeneration of *CDC50* KO dendrites (6) (Figures S3B-S3B”). Surprisingly, this degeneration was completely suppressed in the *orion^1^* hemizygotes (Figures S3C-S3C”), suggesting that the effects of LactC1C2 are mediated by Orion.

To further ask if Orion directly interacts with PS *in vitro*, we purified fat body-derived OrionB-GFP protein from larvae and conducted liposome sedimentation assays (42). As positive and negative controls, we also tested AV-GFP and AV(mut)-GFP (6) which carries mutations that abolish AV/PS interactions (40, 43), respectively. As expected, AV-GFP co-precipitated with PS-containing liposomes but not PC-only liposomes (Figures S3E and S3F). In contrast, AV(mut)-GFP did not co-precipitate with either liposomes (Figures S3E and S3G). Surprisingly, about half OrionB-GFP protein in our assay co-precipitated with PC-only liposomes, and liposome-bound OrionB-GFP increased to 62% when PS was added to the liposomes (Figures S3E and S3H). These data suggest that OrionB may have an intrinsic affinity to phospholipid bilayer and that this interaction is enhanced by the presence of PS.

*In vivo*, we noticed that OrionB-GFP also binds to the surface of several other peripheral tissues, including epidermal cells, muscles, and trachea (Figures 3H-3H”). The OrionB-GFP binding on epidermal cells was enhanced after we gently pinched the larval body wall (Figures S3D-S3D’), which is expected to mildly disrupt the epidermal cell membrane. To understand how much the binding of OrionB-GFP on epidermal cells depends on PS exposure, we overexpressed ATP8A, an ortholog of the PS-specific flippase TAT-1 (9, 44). Overexpression of ATP8A in da neurons is sufficient to suppress PS exposure and the associated dendrite degeneration caused by genetic perturbations of the NAD^+^ pathway (11). The P4-ATPase chaperone CDC50 was co-expressed with ATP8A in a patch of epidermal cells to facilitate ATP8A trafficking. This manipulation drastically suppressed OrionB-GFP binding on the surface of epidermal cells (Figures 3M-3N), suggesting that Orion binding on epidermal cells is largely mediated by PS exposure.

Together, our results suggest that PS exposure induces Orion binding to the cell surface of both neuronal and non-neural cells.

### Orion recruits epidermal Drpr to injured dendrites

Because *drpr* is the only other gene known to be required for epidermal engulfment of PS-exposing dendrites (6, 11), we wondered if Orion and Drpr act in the same pathway. We found that *orion^1^* hemizygous and *drpr* mutant larvae exhibited similar degrees of near complete phagocytosis deficiency in injury-induced dendrite degeneration, as indicated by the portion of unengulfed MApHS-labeled debris (Figures S4A-S4B). We further asked whether removing both *drpr* and *orion* produces stronger phagocytosis defects than the loss of either. We induced whole-body KO of *orion* and *drpr* individually and together. Injured dendrites showed indistinguishable levels of near complete blockage of engulfment in all three genotypes at 24 hrs AI (Figures S4C-S4E), suggesting that *orion* and *drpr* function in the same genetic pathway.

To elucidate the epistatic relationship between *orion* and *drpr*, we tested whether the Orion-PS interaction depends on Drpr and whether gain-of-function (GOF) of one gene can compensate for the loss of the other. At 4-5 hrs AI, OrionB-GFP bound to injured dendrites in wildtype and *drpr* null larvae at similar levels (Figure 4A-4B), demonstrating that Orion does not need Drpr for binding to PS. Two observations further suggest that Orion GOF cannot compensate for the loss of *drpr* in engulfment: First, overexpression of Orion in the fat body did not rescue the engulfment of injured dendrites in *drpr* mutants at 24 hrs AI (Figures S4F and 4C); and second, the degeneration of *CDC50* KO dendrites induced by OrionB-GFP (Figure 3I-3L) was absent in *drpr* mutants (Figures S4G-S4I). To test the effects of *drpr* GOF in *orion* LOF, we overexpressed Drpr in a patch of epidermal cells in the posterior hemisegment (driven by *hh-Gal4*) of *orion^ΔC^* hemizygotes. Interestingly, Drpr OE restored engulfment of injured dendrites, as indicated by the dispersion of dendrite debris specifically in the *hh* domain (Figures 4D-4E). These data suggest that phagocytes with overexpressed Drpr does not absolutely need Orion to detect injured dendrites, which could be due to the ability of Drpr extracellular domain to directly interact with PS (18). However, our data also show that, at physiological levels of Drpr, Orion is indispensable for engulfment.

**Figure 4:**
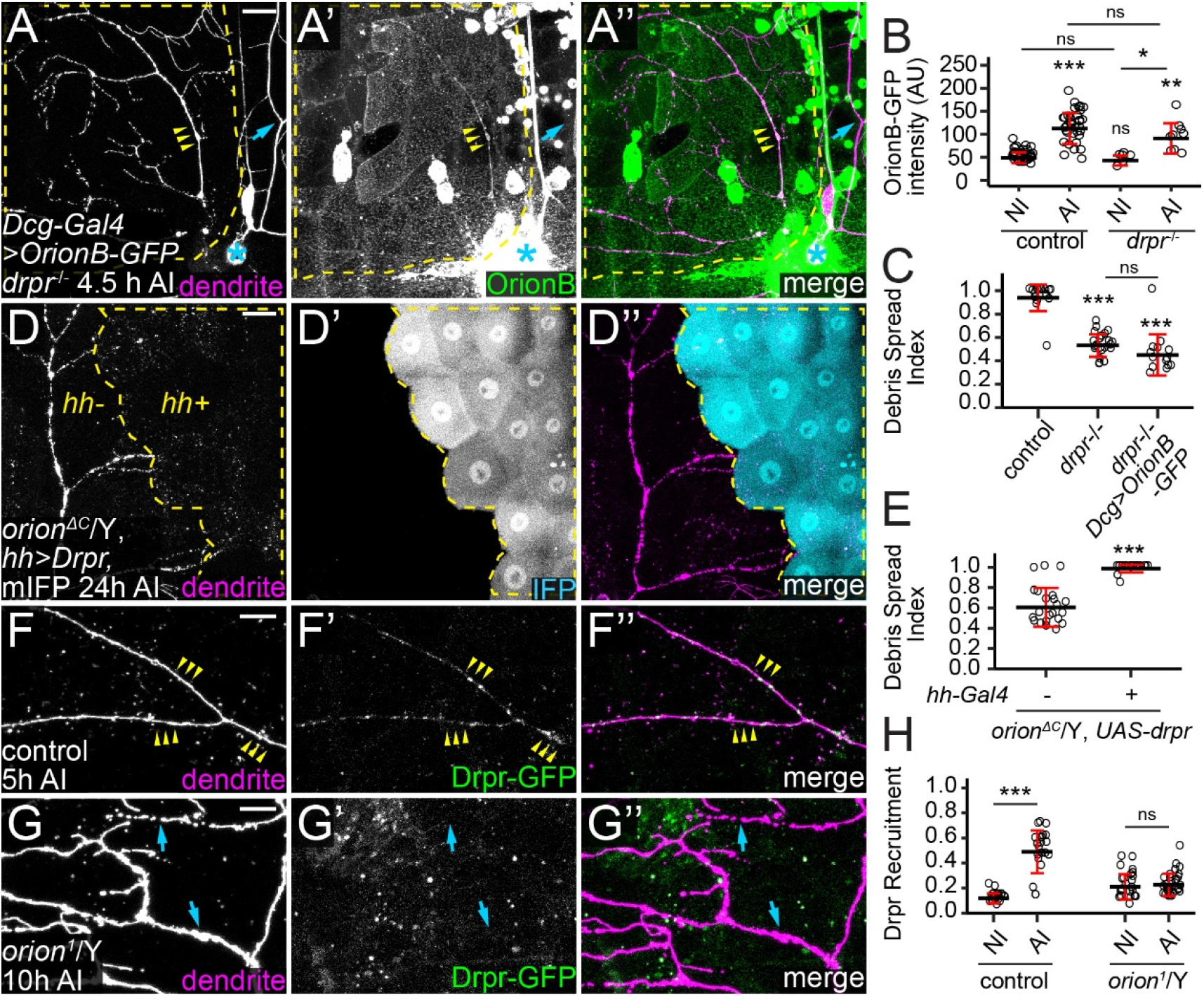
Orion recruits epidermal Drpr to injured dendrites. (A-A”) Labeling of injured dendrites of a ddaC neuron by OrionB-GFP in *drpr*^-/-^ at 4.5 hrs AI. Yellow arrowheads: injured dendrites with OrionB-GFP labeling; blue arrows: uninjured dendrites lacking OrionB-GFP binding. (B) Quantification of Orion-GFP binding on dendrites in the wildtype and the *drpr*^-/-^ larvae. n = number of measurements and N = number of animals: wildtype NI and wildtype AI (same dataset as in Figure 3D); *drpr*^-/-^ NI (n = 6, N = 2); *drpr*^-/-^ AI (n = 9, N = 3). Kruskal-Wallis (One-way ANOVA on ranks) and Dunn’s test, p-values adjusted with the Benjamini-Hochberg method. (C) Quantification of debris spread index of injured dendrites at 22-24 hrs AI. n = number of neurons and N = number of animals: control (same dataset as in 2M); *drpr*^-/-^ (n = 23, N = 10); *drpr*^-/-^ + *Dcg*>*OrionB-GFP* (n = 14, N = 7). One-way ANOVA and Tukey’s test. (D-D”) Engulfment of injured dendrites in an *orion^ΔC^* hemizygous larva with Drpr overexpressed in the *hh* domain. Yellow dash outlines: *hh*>*Drpr, mIFP* region. (E) Quantification of debris spread index of injured dendrites at 22-24 hrs AI in region without Drpr OE and region with Drpr OE in *orion^ΔC^*/Y. n = number of neurons and N = number of animals: region without Drpr OE (n = 22, N = 12); and region with Drpr OE (n = 24, N = 12). Welch’s t-test. (F-F”) Distribution of Drpr-GFP in the presence of injured dendrites in control at 5 hrs AI (F-F”) and in *orion^1^*/Y at 10 hrs AI (G-G”). Yellow arrowheads (F-F”): injured dendrites with Drpr-GFP recruitment; bule arrowheads (G-G”): injured dendrites lacking Drpr-GFP recruitment. (H) Quantification of Drpr-GFP recruitment (Drpr-GFP-positive area on dendrites/total dendrite area). n = measurements: control NI (n = 20, N = 12); control AI (n = 21, N = 12); *orion^1^*/Y NI (n = 27, N = 8); *orion^1^*/Y AI (n = 31, N = 8). Welch’s t-test. For all image panels, neurons were labeled by *ppk-CD4-tdTom*. Yellow dash outlines: territories originally covered by injured dendrites; blue asterisks: injury sites. Scale bars, 25 μm (A-A”, D-D”) and 10 μm (F-G”). For all quantifications, *p≤0.05, **p≤0.01, ***p≤0.001; n.s., not significant. The significance level above each genotype is for comparison with the control. Black bar, mean; red bar, SD. See also Figure S4.

To further understand how *orion* LOF affects Drpr, we examined the distribution of Drpr protein by live imaging. For this purpose, we generated a knock-in allele of *drpr* so that the C-terminus of endogenous Drpr is tagged by mNeonGreen (mNG) (45) (Figure S5J). To further boost the Drpr signal, we also made a Drpr-GFP transgene that contains a genomic fragment of the *drpr* locus tagged with GFP at the Drpr C-terminus (Figure S4J). Combining a copy of each of *drpr-mNG* and *drpr-GFP* (referred to as simply *drpr-GFP* because both fluorescent proteins are green), we were able to see robust Drpr recruitment to injured dendrites prior to dendrite fragmentation (Figures 4F-4F” and 4H), which is consistent with how Drpr functions to instruct phagocytosis.. However, this recruitment was abolished in the *orion^1^* hemizygote, even though injured dendrites showed signs of degeneration, such as thinning and blebbing (Figures 4G-4H). These results suggest that Orion regulates Drpr’s response to degenerating dendrites after injury.

### Orion mediates the interaction between PS and Drpr

Because Orion interacts with PS and functions genetically upstream of Drpr, we wondered whether Orion mediates PS recognition by interacting with Drpr. We first tested whether Orion could bind to Drpr *in vivo* by expressing Drpr in a patch of epidermal cells (driven by *R16D01-Gal4*). Fat body-derived OrionB-GFP was found to accumulate specifically on the surface of these Drpr OE cells in wandering 3^rd^ instar larvae (Figures 5A-5B). Interestingly, some epidermal cells with higher OrionB-GFP enrichment (due to higher Gal4 activities) extruded into neighboring epidermal cells that had low or no OrionB-GFP enrichment (Figures S5A-S5A”). We interpret these extrusions as engulfment of high-OrionB cells by low-OrionB cells. To exclude the possibility that OrionB-GFP was recruited by other cell-surface molecules on epidermal cells as a result of Drpr OE, we also tested a truncated Drpr without the intracellular domain (Drpr^ΔCyto^), which should be defective in intracellular signaling. Overexpression of Drpr^ΔCyto^ also caused drastic OrionB-GFP accumulation on epidermal cells (Figure 5C-5D). These results suggest that OrionB may directly interact with the extracellular domain of Drpr.

**Figure 5:**
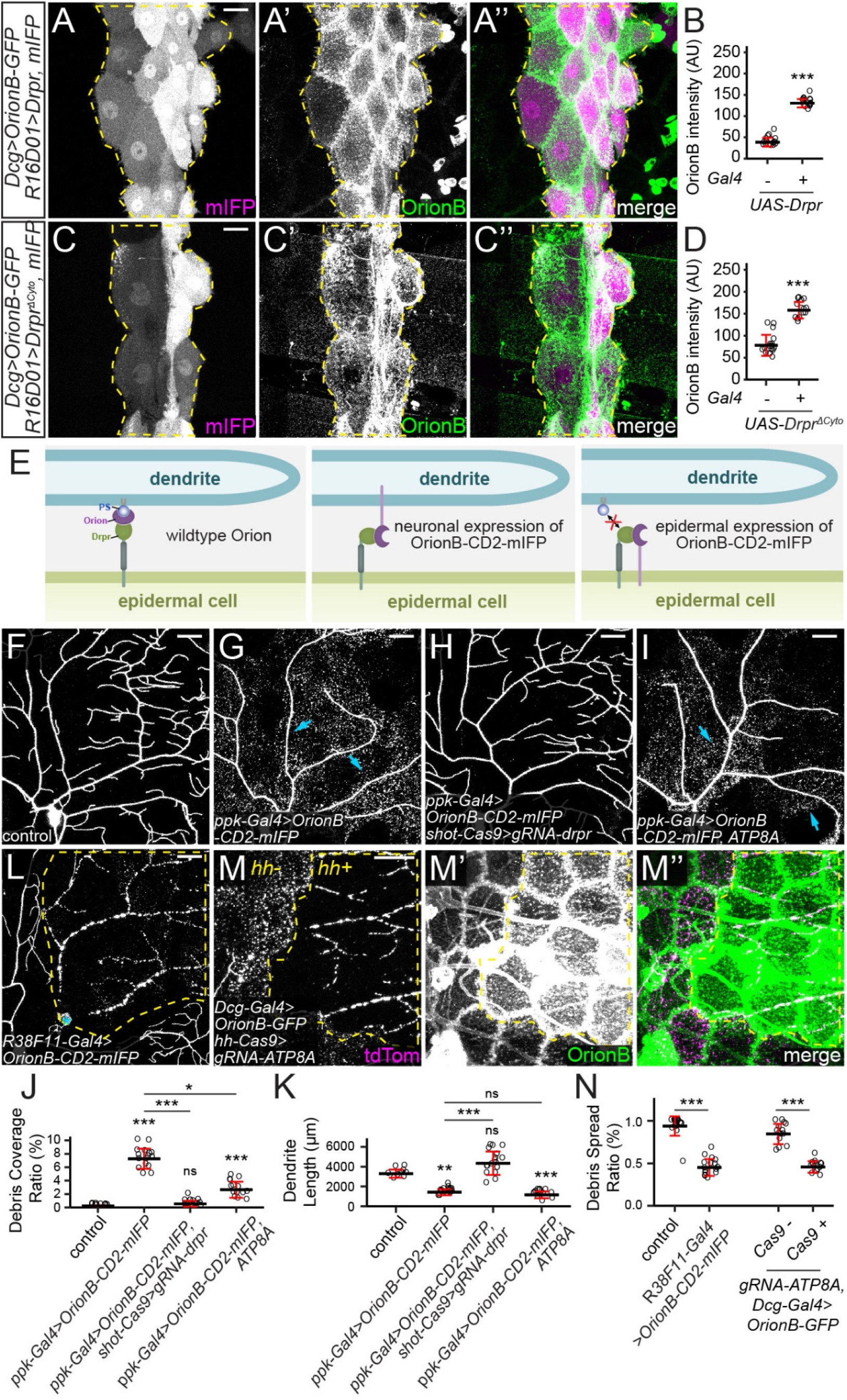
Orion mediates the interaction between PS and Drpr. (A-A”) Distribution of fat body-derived OrionB-GFP with Drpr overexpressed in a patch of epidermal cells driven by R16D01-Gal4. mIFP was coexpressed with Drpr and thus the mIFP intensity is correlated with the Gal4 activity. Yellow dash outlines: Drpr OE region. (B) Quantification of Orion-GFP binding on epidermal cells with or without Drpr OE. n = number of measurements and N = number of animals: without Drpr OE (n = 16, N = 8); with Drpr OE (n = 16, N = 8). Welch’s t-test. (C) Distribution of fat body-drived OrionB-GFP with Drpr^ΔCyto^ overexpressed in a patch of epidermal cells (labeled by mIFP). Yellow dash outlines: Drpr^ΔCyto^ OE region. (D) Quantification of Orion-GFP binding on epidermal cells with or without Drpr^ΔCyto^ OE. n = number of measurements and N = number of animals: without Drpr^ΔCyto^ OE (n = 16, N = 8); with Drpr^ΔCyto^ OE (n = 14, N = 7). Welch’s t-test. (E) Models of Orion function in mediating PS-Drpr interaction (left), interactions between Drpr and Orion-CD2-mIFP overexpressed by neurons (middle) or epidermal cells (right). (F-I) Partial dendritic fields of a control ddaC neuron (F), an OroinB-CD2-mIFP OE neuron (G), an OroinB-CD2-mIFP OE neuron with *drpr* KO in epidermal cells (H), and an OroinB-CD2-mIFP + ATP8A OE neuron (I). (J-K) Quantification of debris coverage ratio (J) and dendrite length (K) at 96 hrs AEL. n = number of neurons and N = number of animals: control (n = 10, N = 6); *ppk-Gal4*>*OroinB-CD2-mIFP* (n = 17, N = 9);*ppk-Gal4*>*OroinB-CD2-mIFP* + *shot-Cas9*>*gRNA-drpr* (n = 17, N = 9);*ppk-Gal4*>*OroinB-CD2-mIFP, ATP8A* (n = 14, N = 7). Kruskal-Wallis (One-way ANOVA on ranks) and Dunn’s test, p-values adjusted with the Benjamini-Hochberg method. (L) Partial dendritic field of an injured ddaC neuron at 25 hrs AI with OroinB-CD2-mIFP expressed in all epidermal cells. (M-M”) Partial dendritic field of an injured ddaC neuron at 9 hrs AI when OroinB-GFP was expressed in fat body and *ATP8A* was knocked out in *hh-* epidermal cells. (N) Quantification of debris spread index of injured dendrites. n = number of neurons and N = number of animals: control (n = 18, N = 10); *R38F11*>*OroinB-CD2-mIFP* (n = 16, N = 8); *Dcg-Gal4*>*OrionB* (n = 14, N = 8); *Dcg-Gal4*>*OrionB* + *hh-Cas9*>*ATP8A* (n = 13, N = 8). Welch’s t-test. Neurons were labeled by *ppk-MApHS* (F-I), and *ppk-CD4-tdTom* (L and M-M”). For all image panels, scale bars, 25 μm. For all quantifications, *p≤0.05, **p≤0.01, ***p≤0.001; n.s., not significant. The significance level above each genotype is for comparison with the control. Black bar, mean; red bar, SD. See also Figure S5.

To further test the hypothesis that Orion mediates the interaction between Drpr and PS, we asked whether expression of an Orion that is permanently tethered to the surface of otherwise wildtype dendrites could bypass the requirement of PS exposure and induce Drpr-dependent phagocytosis. For this purpose, we made an OrionB-CD2-mIFP (mIFP: monomeric infrared fluorescent protein; (46)) transgene in which Orion is located on the extracellular side of the CD2 transmembrane domain (47). As expected, overexpression of OrionB-CD2-mIFP in neurons caused robust dendrite degeneration (Figures 5E-5G, and 5J-5K). We found that this degeneration was completely suppressed by epidermis-specific KO of *drpr* (Figures 5H, and 5J-5K) but was unaffected by suppressing PS exposure in neurons via ATP8A OE (Figures 5I-5K). These data suggest that membrane-tethered Orion is sufficient to induce PS-independent and Drpr-dependent phagocytosis.

Meanwhile, if Orion mediates the recognition of PS by Drpr, we predict that excessive Orion on the surface of epidermal cells would interact with Drpr on the same membrane and interfere with the sensing of PS on dendrites. We first tested this idea by overexpressing OrionB-CD2-mIFP in all epidermal cells (Figure 5E). Indeed, this manipulation fully blocked the engulfment of injured dendrites at 25 hrs AI (Figures 5L and 5N). Drpr was robustly detected on the cell membranes of OrionB-CD2-mIFP-expressing cells (Figures S5C and S5D), suggesting that the impaired engulfment was not due to defects in Drpr subcellular localization. We then tested whether accumulation of secreted Orion on the surface of epidermal cells has a similar effect in blocking engulfment. Consistent with the idea that Orion binds PS, *ATP8A* KO in epidermal cells resulted in a drastically increased surface level of OrionB-GFP (Figure 5M’). This OrionB-GFP accumulation was associated with phagocytosis deficiency, as indicated by the lack of spread of dendritic debris at 9 hrs AI specifically in the *ATP8A* KO cells (Figure 5M-5N).

Together, our results show that interactions between Orion and Drpr from the same versus apposing membranes produce opposite phenotypes (defective versus dominant dendrite engulfment, respectively) and support the idea that Orion functionally bridges PS and Drpr.

### The Orion dosage determines the sensitivity of epidermal cells to PS-exposing dendrites

While examining the phenotypes of the *orion^1^* mutant, we noticed impaired phagocytosis in *orion^1^*/+ heterozygous larvae: Neurons with *CDC50* KO and TMEM16F OE showed no signs of membrane loss or degeneration in these animals (Figures 6A-6C, 6J-6K), comparable to those in *orion^1^* hemizygotes. However, the engulfment of injured dendrites was normal in *orion^1^*/+ (Figures S6A-S6B). Considering that *CDC50* KO and TMEM16F OE causes milder PS exposure than does injury (6), these data suggest that removing half the dosage of Orion reduces the sensitivity of epidermal cells to PS exposure on dendrites but does not block phagocytosis when PS exposure is high. To further test if the Orion dosage determines the ability of epidermal cells to sense PS-exposing dendrites, we increased the Orion dosage by adding an *orion* duplication (Dp(1;3)DC496) to the wildtype. The *orion* duplication strongly enhanced the debris level and dendrite loss in a *CDC50* KO background (Figures 6E, 6J-6K), which by itself only causes weak PS exposure (6) and low levels of debris (Figures 6D, 6J-6K). Thus, extra Orion can indeed increase the sensitivity of epidermal cells to neuronal PS.

**Figure 6:**
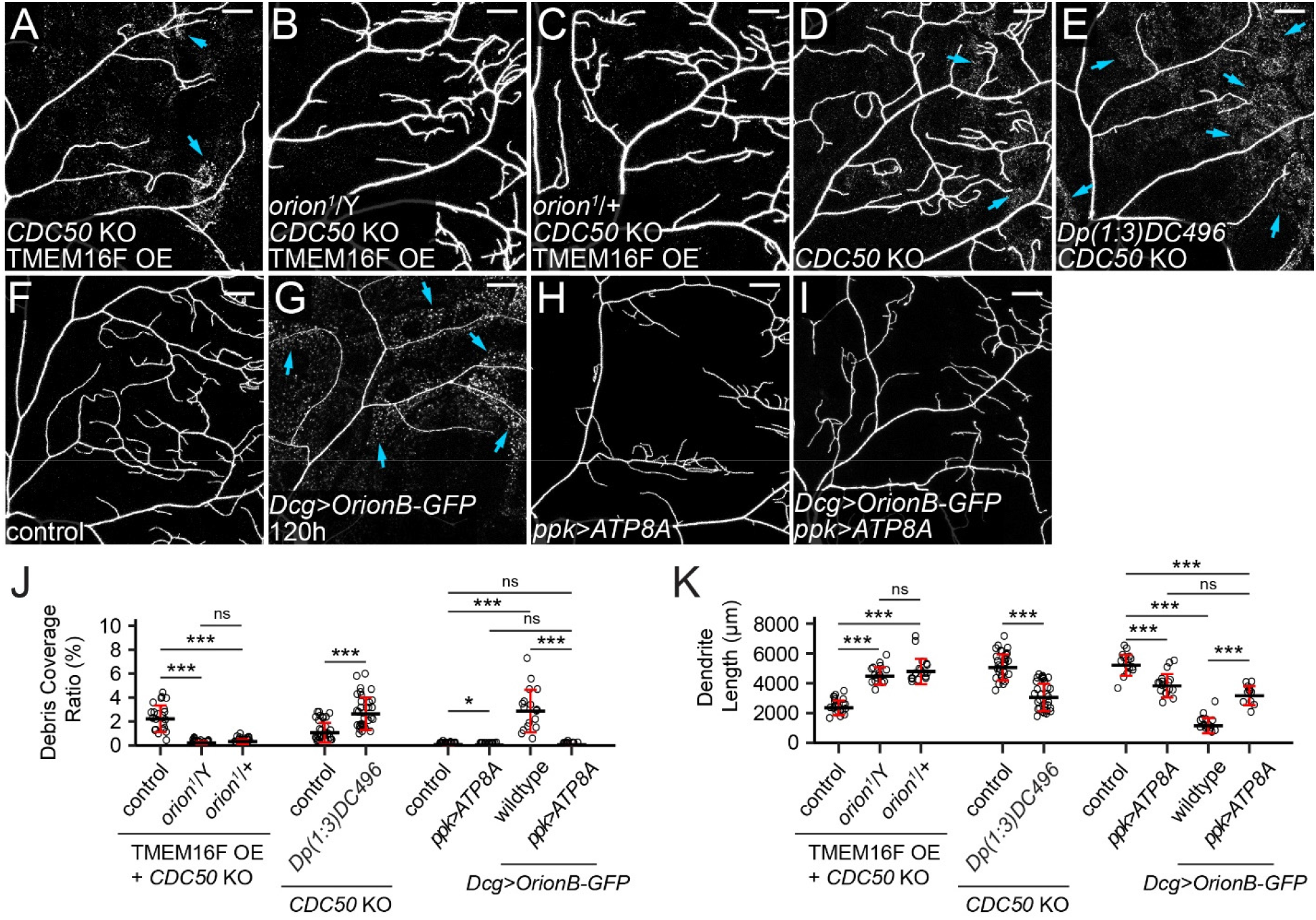
The Orion dosage determines the sensitivity of epidermal cells to PS-exposing dendrites. (A-C) Partial dendritic fields of a TMEM16F OE +*CDC50* KO ddaC neuron in the WT background (A, same image as Figure 1E), in the *orion^1^* hemizygous background (B, same image as Figure 1F), and in the *orion^1^* heterozygous background (C). (D-E) Partial dendritic fields of *CDC50* KO neurons in the control (D) and Dp(1;3)DC496 (E) at 120 hrs AEL. Blue arrows: debris shed from dendrites. (F-I) Partial dendritic fields of ddaC neurons in the control (F), with fat body-derived OrionB-GFP (G), with ATP8A OE in the neuron (H), and with fat body-derived OrionB-GFP and ATP8A OE in the neuron (I) at 120 hrs AEL. (J-K) Quantification of debris coverage ratio (J) and dendrite length (K). n = number of neurons and N = number of animals: for TMEM16F OE +*CDC50* KO, control and *orion^1^*/Y (same dataset as in Figure 1G), *orion^1^*/+ (n = 18, N = 9); for (J), Kruskal-Wallis (One-way ANOVA on ranks) and Dunn’s test, p-values adjusted with the Benjamini-Hochberg method; for (K), one-way ANOVA and Tukey’s test. For *CDC50* KO, control (n = 33, N = 17), Dp(1;3)DC496 (n = 33, N = 17), Welch’s t-test. For effects of *Dcg*>*OrionB-GFP* and *ppk*>*ATP8A* at 120 hrs AEL, control (n = 17, N = 9),*ppk*>*ATP8A* (n = 17, N = 9), *Dcg*>*OrionB-GFP* (n = 17, N = 9),*ppk*>*ATP8A* + *Dcg*>*OrionB-GFP* (n = 12, N = 7); for (J), Kruskal-Wallis (One-way ANOVA on ranks) and Dunn’s test, p-values adjusted with the Benjamini-Hochberg method; for (K), one-way ANOVA and Tukey’s test. Neurons were labeled by *ppk-MApHS* (A-C and H), *ppk-CD4-tdTom* (D-G), and *ppk-Gal4*>*CD4-tdTom* (I). For all image panels, scale bars, 25 μm. For all quantifications, *p≤0.05, ***p≤0.001; n.s., not significant. The significance level above each genotype is for comparison with the control. Black bar, mean; red bars, SD. See also Figure S6.

Further supporting this notion, we found that fat body-derived OrionB bound to wildtype dendrites at 120 hrs AEL (Figures S6C) and induced pronounced degeneration (Figures 6F-6G and 6J-6K). This degeneration is PS-dependent because it was completely suppressed by ATP8A OE in neurons (Figure 6H-6I, and 6J-6K). This surprising result demonstrates that, in late larval stages, wildtype dendrites expose low levels of PS that cannot be detected by common PS sensors such as AV and LactC1C2, but can be bound by overexpressed Orion.

### CX_3_C and RRY motifs are important for Orion function

Orion shares a CX3C motif with the human chemokine CX3CL1 and an RRY motif with several human neutrophil peptides. The CX3C motif is required for Orion’s function in MB remodeling (24), but the importance of the RRY motif in Orion has not been investigated. To test whether these motifs play any role in PS-mediated dendrite engulfment, we generated *UAS-OrionB-GFP* variants carrying mutations in them (OrionB^AX3C^ and OrionB^AAY^). We first compared these OrionB-GFP variants in their abilities to potentiate dendrite degeneration of *CDC50* KO neurons, which would be reflected by increased debris level and reduction of dendrite length. Although fat body-derived OrionB^AX3C^ and OrionB^AAY^ both potentiated the debris level of *CDC50* KO neurons (Figures 7A-7D, and 7I), only OrionB^AX3C^, but not OrionB^AAY^, caused a weak (21%) dendrite reduction of *CDC50* KO neurons (Figure 7J). In comparison, WT OrionB caused 85% reduction of *CDC50* KO dendrites (Figure 7J). We next tested OrionB-GFP variants in rescuing *orion^ΔC^* hemizygotes with the dendrite injury assay. Orion^AX3C^ restored the engulfment of injured dendrites to the same level as WT OrionB-GFP while OrionB^AAY^ only partially restored the engulfment (Figures 7E-7H, and 7I). These results suggest that these mutations impair, but do not abolish, Orion activity.

**Figure 7:**
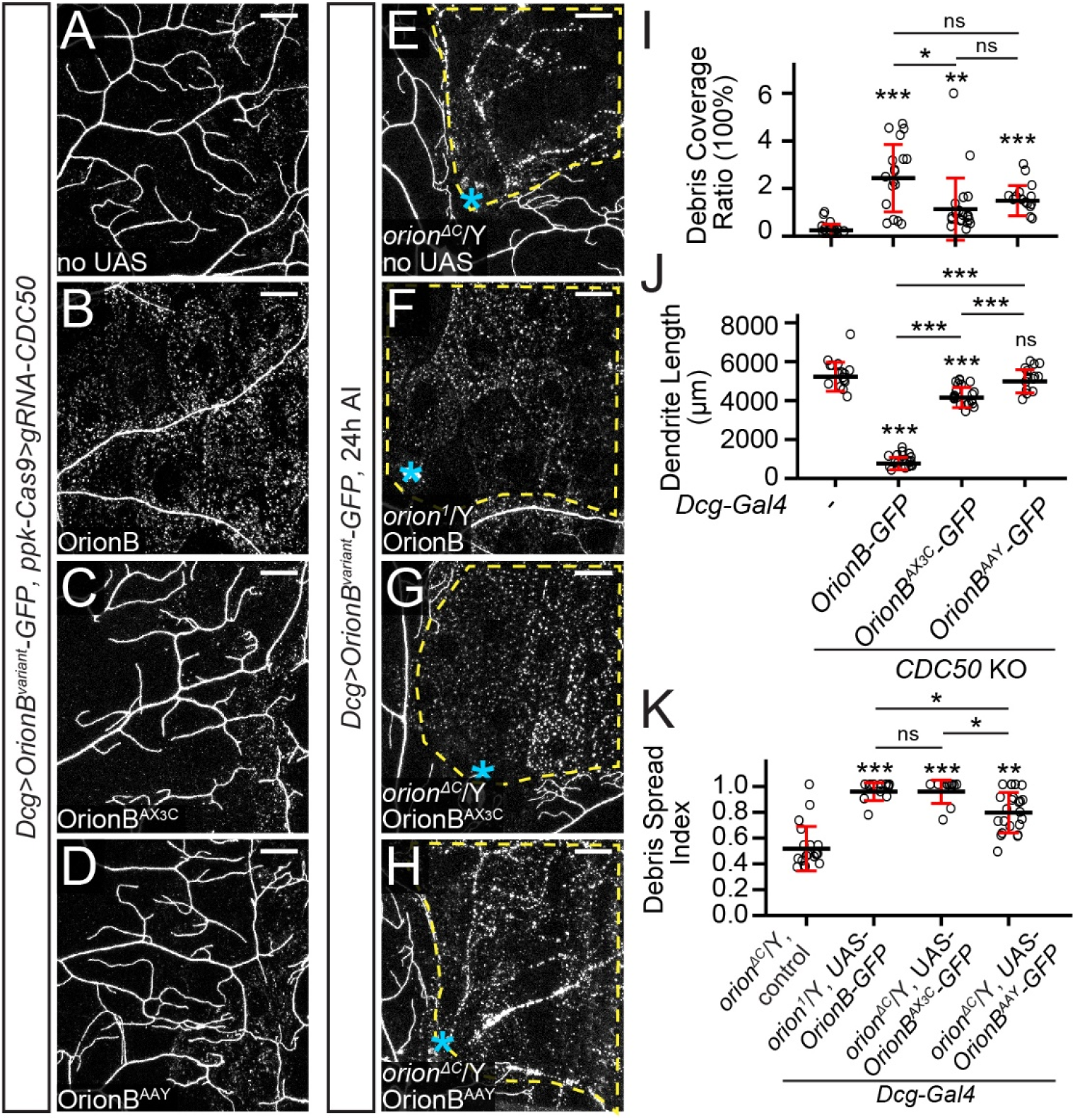
CX_3_C and RRY motifs are important for Orion function. (A-D) Partial dendritic fields of *CDC5O* KO ddaC neurons without OrionB variants (A), with WT OrionB=GFP (B), OrionB^AX3C^-GFP (C), and OrionB^AAY^-GFP (D) expressed in the fat body at 120 hrs AEL. (I-J) Quantification of debris coverage ratio (I) and dendrite length (J) at 120 hrs AEL. n = number of neurons and N = number of animals: *CDC50* KO control (n = 16, N = 8); *CDC50* KO + *Dcg*>*OrionB-GFP* (n = 18, N = 9); *CDC50* KO + *Dcg*>*OrionB^AX3C^-GFP* (n = 20, N = 10); *CDC50* KO + *Dcg*>*OrionB^AAY^-GFP* (n = 16, N = 8); *CDC50* KO +*Dcg*>*OrionB^1^-GFP* (n = 15, N = 8). For (I), Kruskal-Wallis (One-way ANOVA on ranks) and Dunn’s test, p-values adjusted with the Benjamini-Hochberg method; for (J), one-way ANOVA and Tukey’s test. (E-H) Partial dendritic fields of ddaC neurons in *orion^ΔC^* or *orion^1^* hemizygous larvae without OrionB variants (E), with WT OrionB=GFP (F), OrionB^AX3C^-GFP (G), and OrionB^AAY^-GFP (H) expressed in the fat body at 22-24 hrs AI. Yellow dash outlines: territories originally covered by injured dendrites; blue asterisks: injury sites. (K) Quantification of debris spread index of injured dendrites. n = number of neurons and N = number of animals: *orion^ΔC^*/Y control (n = 19, N = 9); *orion^1^*/Y *+* *Dcg*>*OrionB-GFP* (same dataset as in 2M); *orion^ΔC^*/Y + *Dcg*>*OrionB^AX3C^-GFP* (n = 12, N = 7); *orion^ΔC^*/Y + *Dcg*>*OrionB^AAY^-GFP* (n = 23, N = 12); *orion^ΔC^*/Y + *Dcg*>*OrionB^1^-GFP* (n = 15, N = 8). Kruskal-Wallis (One-way ANOVA on ranks) and Dunn’s test, p-values adjusted with the Benjamini-Hochberg method. For all image panels, neurons were labeled by *ppk-CD4-tdTom*. Scale bars, 25 μm. For all quantifications, *p≤0.05, **p≤0.01, ***p≤0.001; n.s., not significant. The significance level above each genotype is for comparison with the control. Black bar, mean; red bars, SD.

## DISCUSSION

### Orion mediates interactions between PS and Drpr

By triggering phagocytosis, the recognition of PS exposed on neurons is a critical event during neurodegeneration and clearance. Although several studies have implied the involvement of Drpr in this process in *Drosophila* (6, 11, 15, 19–21, 28), how Drpr mediates PS recognition *in vivo* is unclear. In this study, we present several lines of *in vivo* evidence that strongly indicate that the *Drosophila* chemokine-like Orion is a PS-binding bridging molecule that enables Drpr to respond to neuronal PS exposure. First, Orion is required for multiple scenarios of Drpr-dependent phagocytosis of sensory dendrites and functions upstream of Drpr. Second, Orion binds to PS-exposing cell surfaces. We show that Orion binds to neurons and epidermal cells that expose PS as a result of tissue-specific KO of the PS flippase ATP8A. In addition, ATP8A OE, which retains PS in the inner membrane leaflet, eliminates Orion binding on epidermal cells and late-larval dendrites, suggesting that this binding is PS-dependent. Importantly, Orion outcompetes Annexin V for binding to injured dendrites, suggesting that Orion may directly interact with PS *in vivo*. Third, overexpressed Drpr proteins can trap Orion on the cell surface, suggesting that Drpr interacts with Orion *in vivo*. Lastly, when expressed in neurons, membrane-tethered Orion bypasses the requirement for PS in inducing Drpr-dependent engulfment, but when expressed in phagocytes, membrane-tethered Orion blocks PS-induced engulfment. Based on these observations, we propose that Orion functions as a bridging molecule between PS and Drpr.

Previously, SIMU, a PS-binding transmembrane protein expressed by *Drosophila* embryonic phagocytes, was proposed to be a bridging molecule (48, 49). Orion and SIMU contribute to phagocytosis through distinct mechanisms. First, SIMU is expressed by phagocytes to allow them to tether apoptotic cells (48), while Orion is secreted from many peripheral tissues and functions as an opsonin to enable phagocytosis. Second, SIMU is a membrane protein that shares homology with Drpr but functions at a different step in apoptotic neuron clearance compared to Drpr (48). In contrast, as a secreted protein, Orion functions at the same step of phagocytosis as Drpr. Therefore, SIMU behaves more like a tethering receptor (49), while Orion is functionally analogous to PS-bridging molecules in other species (41, 50). Although we focus our analyses on the engulfment of somatosensory dendrites, the ubiquitous roles of PS and Drpr in phagocytosis and the broad expression patterns of Orion suggest that Orion may be widely involved in PS-mediated phagocytosis in *Drosophila*. This view is supported by our findings that Orion deposited in the hemolymph can mediate phagocytosis in distant tissues and that the accumulation of Orion on epidermal cells turns these cells into targets of phagocytosis.

### The level of Orion modulates phagocyte sensitivity to PS

Although the role of PS exposure in inducing phagocytosis has been well documented (7, 51), what determines the sensitivity of phagocytes to PS is much less understood. In this study, we discovered that the available level of Orion is a determinant of phagocyte sensitivity to PS in *Drosophila*. We show that reducing the dosage of functional Orion by one half makes phagocytes blind to dendrites with moderate levels of PS exposure (i.e. *CDC50* KO + TMEME16F OE neurons), but the reduced Orion does not affect the ability of phagocytes to engulf dendrites that display high levels of PS exposure (i.e. injury). Conversely, an extra copy of the *orion* locus enhances the ability of phagocytes to engulf dendrites that have mild PS exposure (i.e. *CDC50* KO). These results suggest that endogenous Orion is likely expressed at a balanced level to enable the right amount of phagocytosis: Too much Orion may cause unintended phagocytosis of stressed cells that display mild PS exposure, while too little Orion may interfere with efficient clearance of sick cells or structures that are beyond rescue. Consistent with this idea, endogenous Orion is expressed at a low level during larval development (Figure S2G and S2I), but is dramatically upregulated during metamorphosis (24), a time when large-scale tissue remodeling and clearance take place (52).

### Orion has distinct roles in neurite maintenance and remodeling

Orion was previously known to be required for axonal pruning and clearance of MB **γ** Kenyon neurons during metamorphosis (24). In that context, Orion is expressed in the remodeling MB neurons and functions as a “find-me” signal for glia to penetrate the axon bundles and engulf axonal debris. In contrast, in the larval peripheral nervous system (PNS), Orion is supplied by many non-neural tissues and functions as a permissive signal for phagocytosis of sick or broken dendrites. This distinction is likely due to two differences between the larval PNS and the remodeling CNS. First, in the PNS, the dendrites of da neurons are exposed to the hemolymph and are readily accessible to Orion that is secreted from other tissues, whereas in the CNS, the axons are more tightly packed and may be harder for extrinsic Orion to access. Neuron-derived Orion would thus be more effective than extrinsic Orion for promoting phagocytosis in the CNS. Intriguingly, we detected Orion expression in a small number of neurons in the larval ventral nerve cord (VNC). It will be interesting to find out whether these neurons are particularly subject to degeneration. Second, compared to the larval PNS where degenerative events are rare, the nervous system undergoing metamorphosis has a much greater demand for phagocytosis. Turning on Orion expression in neurons may thus be required for efficient clearance of all pruned neurites. Consistent with this idea, we detected Orion expression also in a subset of da neurons during metamorphosis.

### Orion possesses unique properties compared to other PS-binding molecules

Fluorescent PS probes based on AV and Lact are widely used to visualize PS exposure in cell culture and live animals and have been crucial for many discoveries in PS biology (6, 40, 41, 53, 54). LactC1C2 is known to have a higher affinity to PS than AV (55–57). We previously observed additional differences between these two proteins. Compared to LactC1C2, which coats the dendrite surface well, AV tends to traffic to endocytic vesicles inside PS-exposing dendrites, consistent with the ability of the AV complex to induce endocytosis upon PS-binding (58, 59). Importantly, unlike AV, which does not alter the kinetics of neurite degeneration, LactC1C2 binding to PS-exposing dendrites potentiates Drpr-dependent degeneration (6). This latter observation previously led us to hypothesize that LactC1C2 may contain unknown sequences that interact with Drpr (6). In this study, we show that the effect of LactC1C2 in exacerbating dendrite degeneration depends on Orion, suggesting instead that LactC1C2 may indirectly promote phagocyte/dendrite interactions by enhancing Orion function. One possible mechanism is that LactC1C2 binding on the plasma membrane further disrupts the membrane and causes more PS exposure that can subsequently be detected by endogenous Orion.

Compared to AV and LactC1C2, Orion displays unique lipid binding properties. Our *in vivo* evidence suggests that Orion may have a higher affinity for PS than AV, as Orion efficiently outcompetes AV in binding to injured dendrites. In addition, overexpressed Orion binds to healthy epidermal cells and dendrites (albeit the latter only in wandering 3^rd^ instar larvae), while AV and LactC1C2 do not (6). Although our *in vitro* evidence suggests that Orion may have an intrinsic affinity for phospholipid bilayers, *in vivo* Orion-binding to epidermal cells and WT dendrites largely depends on PS exposure. One possibility is that other endogenous factors modify Orion-lipid interactions and make Orion more specific to PS *in vivo*.

The surprising finding that overexpressed Orion binds to peripheral tissues and dendrites suggest that these cells may expose PS under physiological conditions, perhaps at a level too low to detect by AV and LactC1C2. Thus, low levels of PS exposure may be much more prevalent *in vivo* than previously thought. Intriguingly, binding of overexpressed Orion induces degeneration of wildtype dendrites but does not cause obvious phagocytosis of other non-neural tissues, suggesting that neurons may be more vulnerable to PS-induced phagocytosis than other cell types.

### Functional conservation between Orion and human immunomodulatory proteins

Recently, many human chemokines were found to bind to PS exposed on apoptotic vesicles and serve as “find-me” signals to attract phagocytes (60). Orion shares the CX_3_C motif with the mammalian chemokine CXC3L1 and has also three glycosaminoglycan (GAG) putative binding sequences, a hallmark of chemokine activity (61). Even though direct binding to PS has not been demonstrated for CXC3L1, this chemokine is required for microglia-mediated synapse elimination after whisker lesioning (25), a process likely involving PS exposure (62). In addition, Orion contains a RRY motif commonly found in human neutrophil peptides, small antimicrobial peptides important for innate immunity (63). We found that both CX_3_C and RRY motifs are important for Orion function in mediating phagocytosis of neurons. Although Orion does not show global sequence homologies to mammalian immunomodulatory proteins, its interaction partner Drpr has mammalian homologs that are involved in phagocytosis of neurons through unknown mechanisms (17). Thus, the common features between Orion and human proteins indicate that a functional conservation may exist between PS-sensing mechanisms in insects and humans.

## METHODS

### Fly strains

The details of fly strains used in this study are listed in Table S1 (Key Resource Table). For labeling of C4da neurons, we used *ppk-MApHS*, *ppk-CD4-tdTom*, and *ppk-Gal4 UAS-CD4-tdTom*. For labeling of all da neurons, we used *21-7-Gal4 UAS-CD4-tdTom*. For labeling PS exposure on dendrites, we used *dcg-Gal4 UAS-AnnexinV-mCard* and *dcg-Gal4 UAS-GFP-LactC1C2*. For visualizing OrionB labeling on cell surface, we used *dcg-Gal4 UAS-OrionB-GFP*, *Dcg-LexA LexAop-OrionB-GFP*, and *R16A03-LexA LexAop-OrionB-GFP*. We generated *dcg-LexA* by converting the Gal4 in *dcg-Gal4* into LexAGAD. The conversion process will be published elsewhere.

See Supplemental Methods for details of molecular cloning and transgenic flies, generation of KI flies, CRISPR-TRiM, live imaging, immunohistochemistry, protein purification, liposome sedimentation assay, image analysis and quantification, and statistical analysis. See Table S2 for gRNA target sequences.

## DATA SHARING PLANS

All study data are included in the article and/or Supplemental Methods. Constructs and fly strains will be available upon request.

## ACKNOWLEDGMENTS

We thank Larry Zipursky, Loren Looger, Xinhua Lin, BACPAC Resources Center, *Drosophila* Genomics Resource Center (DGRC), the National Cancer Institute (NCI), and Addgene for plasmids; Marc Freeman and Bloomington Stock Center for fly stocks; Cornell BRC Imaging facility for access to microscopes (funded by NIH grant S10OD018516); Cornell CSCU for advice on statistics; lab members of Chris Fromme, Jeremy Baskin, and Yuxin Mao for technical assistance and advice on biochemistry; Shaogeng Tang for experimental tests; Mike Goldberg for critical reading and suggestions on the manuscript. This work was supported by NIH grants (R01NS099125 and R21OD023824) awarded to C.H., and by ARC grant (PJA 20151203422) and FRM grant (DEQ20160334870) awarded to J.-M.D.

## FIGURES AND FIGURE LEGENDS

**Figure S1:**
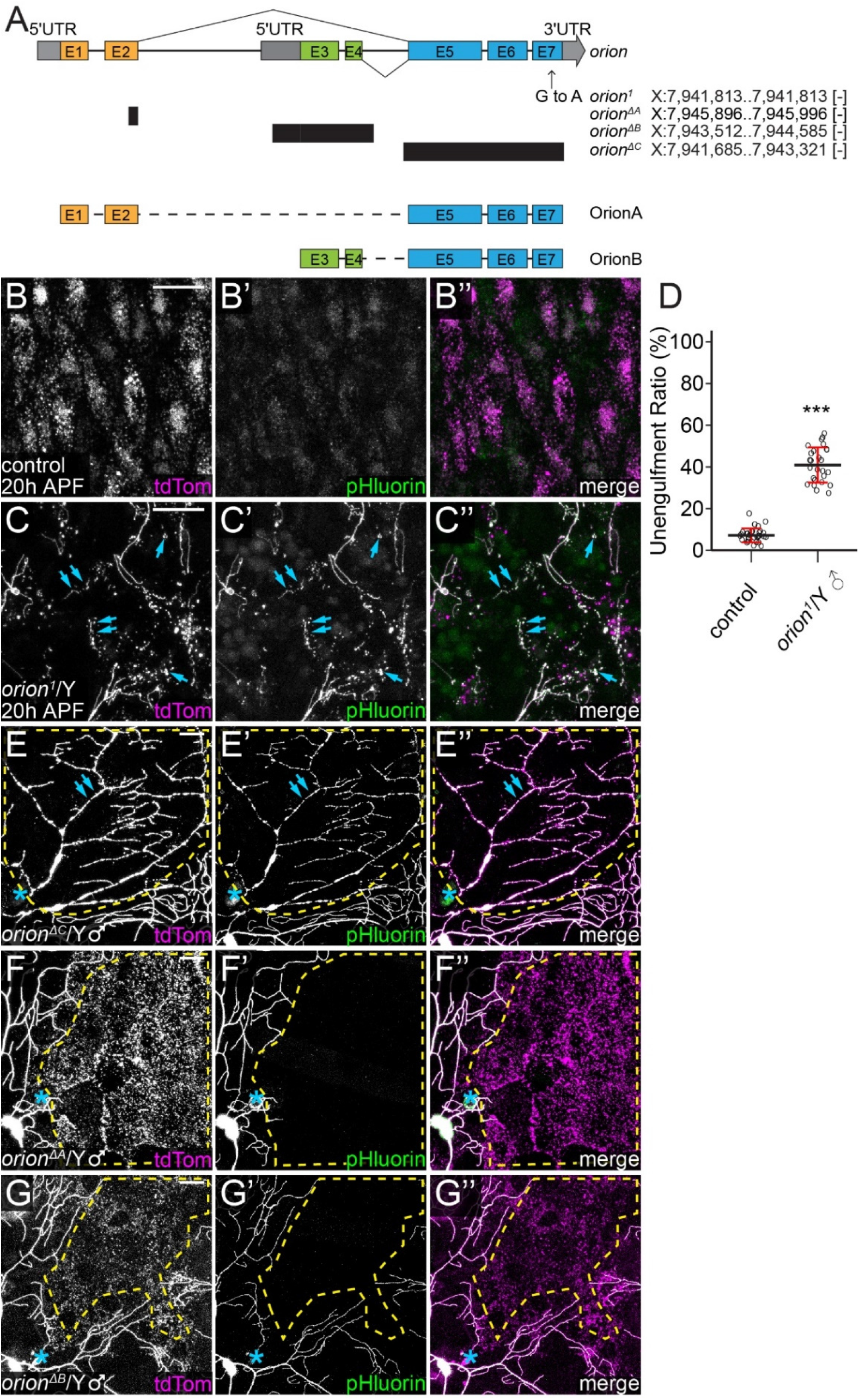
*orion* mutant alleles and engulfment phenotypes. (A) Schematic representation of the *orion* locus in the wildtype and in four different *orion* alleles (*orion^1^, orion^ΔA^, orion^ΔB^* and *orion^ΔC^*). Black bars indicate deleted regions. The two mRNA splicing isoforms of *orion* are also indicated. The *orion* gene annotation is based on Release 6 reference genome assembly. (B-C”) Partial dendritic fields of ddaC neurons at 20 hrs after puparium formation (APF) in wildtype (B-B”) and *orion^1^* hemizygous (C-C”) pupae. Blue arrows: unengulfed dendrite fragments. (D) Quantification of unengulfment ratio of pruned dendrites (area of pHluorin-positive debris/area of tdTom-positive debris) at 19-21 hrs APF. n = number of neurons and N = number of animals: control (n = 30, N = 12); *orion^1^* hemizygotes (n = 26, N = 9). Welch’s t-test; ***p≤0.001; black bar, mean; red bars, SD. (E-E”) Partial dendritic field of a ddaC neuron in *orion^ΔC^* hemizygotes at 23 hrs AI. Blue arrows: injured but unengulfed dendrite fragments. (F-G”) Partial dendritic fields of ddaC neurons in *orion^ΔA^* hemizygotes (G-G”), and *orion^ΔB^* hemizygotes (G-G”) at 23-24 hrs AI. In all image panels, neurons were labeled by *ppk-MApHS*. Scale bars, 25 μm. In (E-G”), yellow dash outlines: territories originally covered by injured dendrites; blue asterisks: injury sites.

**Figure S2:**
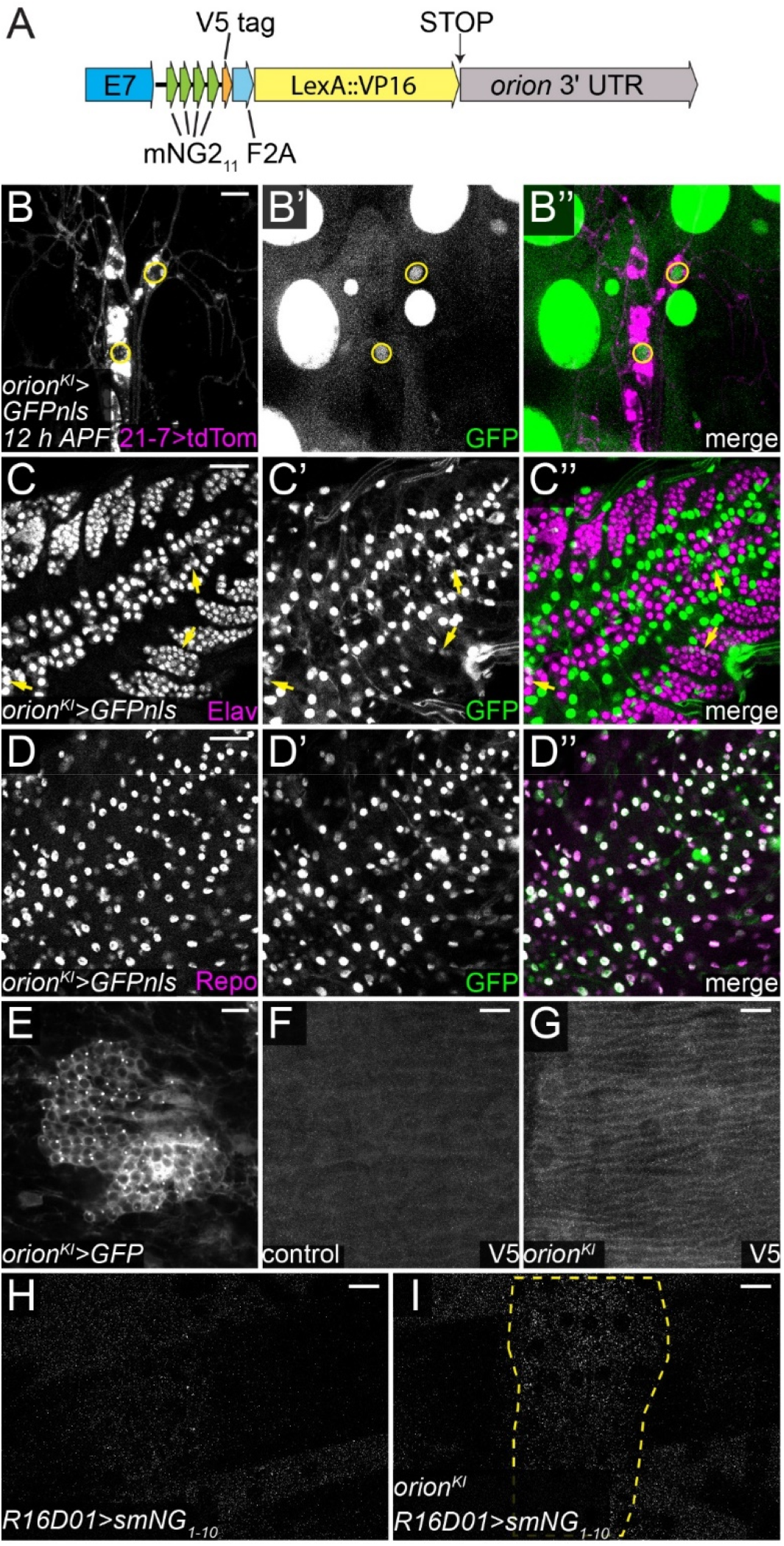
Orion functions cell-non-autonomously. (A) Schematic representation of the *orion^KI^* allele. (B-B”) GFPnls expression driven by *orion-LexA* in ddaC and ddaE neurons at 12 hrs APF. Yellow circles indicate nuclei of ddaC and ddaE neurons. Neurons were labeled by *21-7-Gal4*>*CD4-tdTom*. (C-D”) GFPnls expression driven by *orion-LexA* in the larval ventral nerve cord (VNC). Nuclei of neurons are labeled by Elav staining (C-C”); glial nuclei are labeled by Repo staining (D-D”). Yellow arrows (C-C”): a small number of neurons expressing *orion*. (E) GFP expression driven by *orion-LexA* in the mushroom body Kenyon cells at 8 hrs APF. (F-G) V5 staining of larval body walls in the wildtype control (*w^1118^*) (F) and *orion^KI^* (G). (H-I) Signals of reconstituted mNG in control (H) and *orion^KI^* (I) epidermal cells. Secreted mNG1-10 (smNG1-10) was expressed in patches of epidermal cells (outlined in H) driven by *R16D01-Gal4*. Scale bars, 10 μm (B and E) and 25 μm (C-D” and F-I).

**Figure S3:**
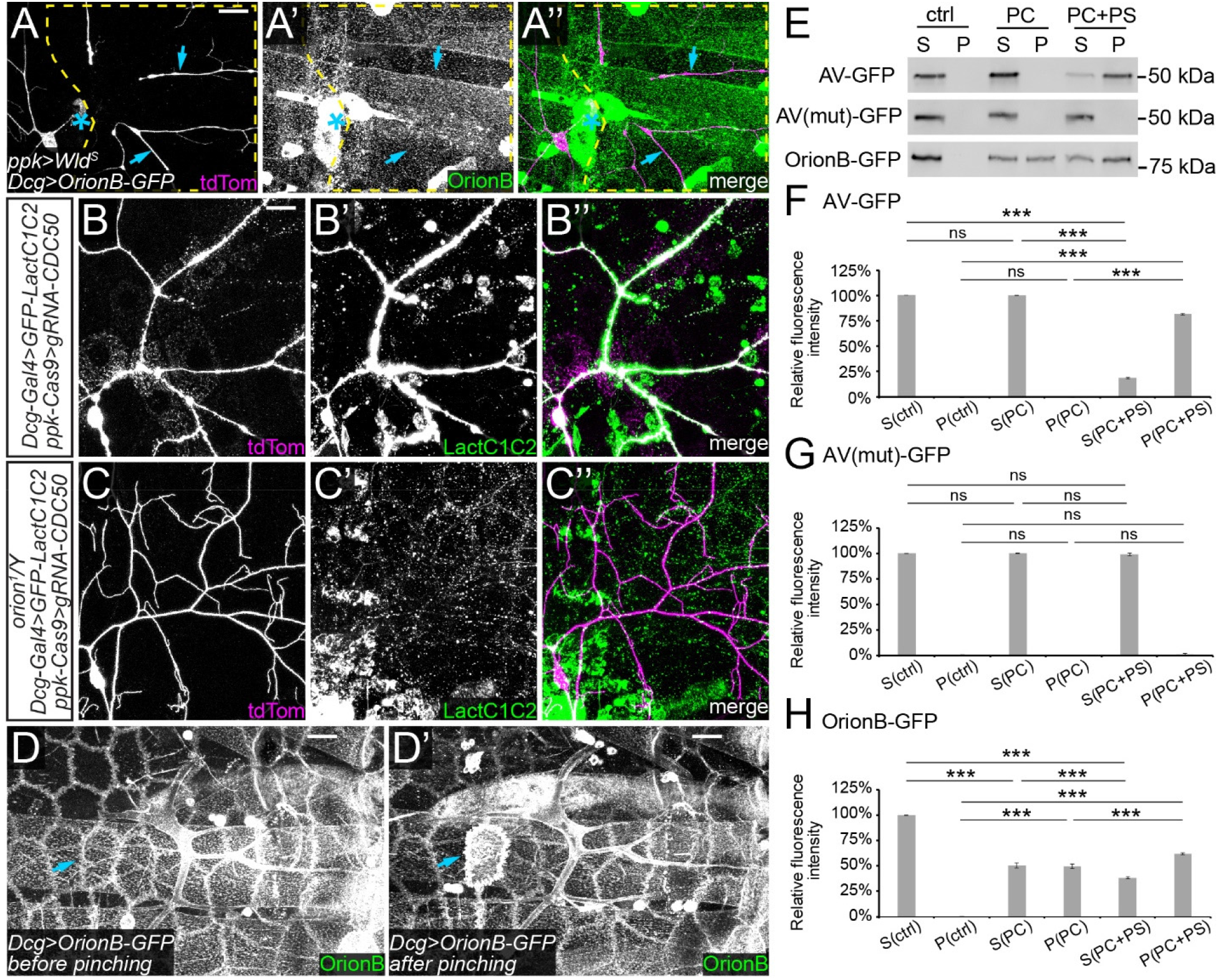
PS exposure induces Orion binding to the cell surface. (A-A”) Lack of OrionB-GFP labeling on injured dendrites of a ddaC neuron with Wld^S^ OE at 6 hrs AI. Yellow dash outlines: territories originally covered by injured dendrites; blue asterisks: injury sites; blue arrows: injured dendrites lacking OrionB-GFP labeling. (B-C”) *CDC50* KO neurons in the presence of fat body-derived GFP-LactC1C2 in control (B-B”) and *orion^1^* hemizygous (C-C”) larvae. (D-D’) OrionB-GFP binding on epidermal cells before (D) and after (D’) gentle pinching. Blue arrows indicate enhanced OrionB binding. (E) Western blot of liposome sedimentation assay. Ctrl, control without liposomes; PC, PC-only liposomes; PC+PS, liposomes containing 20% PS; S, supernatant; P, pellet. (F-H) Quantification of signal intensity of AV-GFP (F), AVmut-GFP (G), and OrionB-GFP (H) in the western blot of liposome sedimentation assay. n=3. Neurons were labeled by *ppk-CD4-tdTom* (A-C”). In all image panels, scale bars, 25 μm.

**Figure S4:**
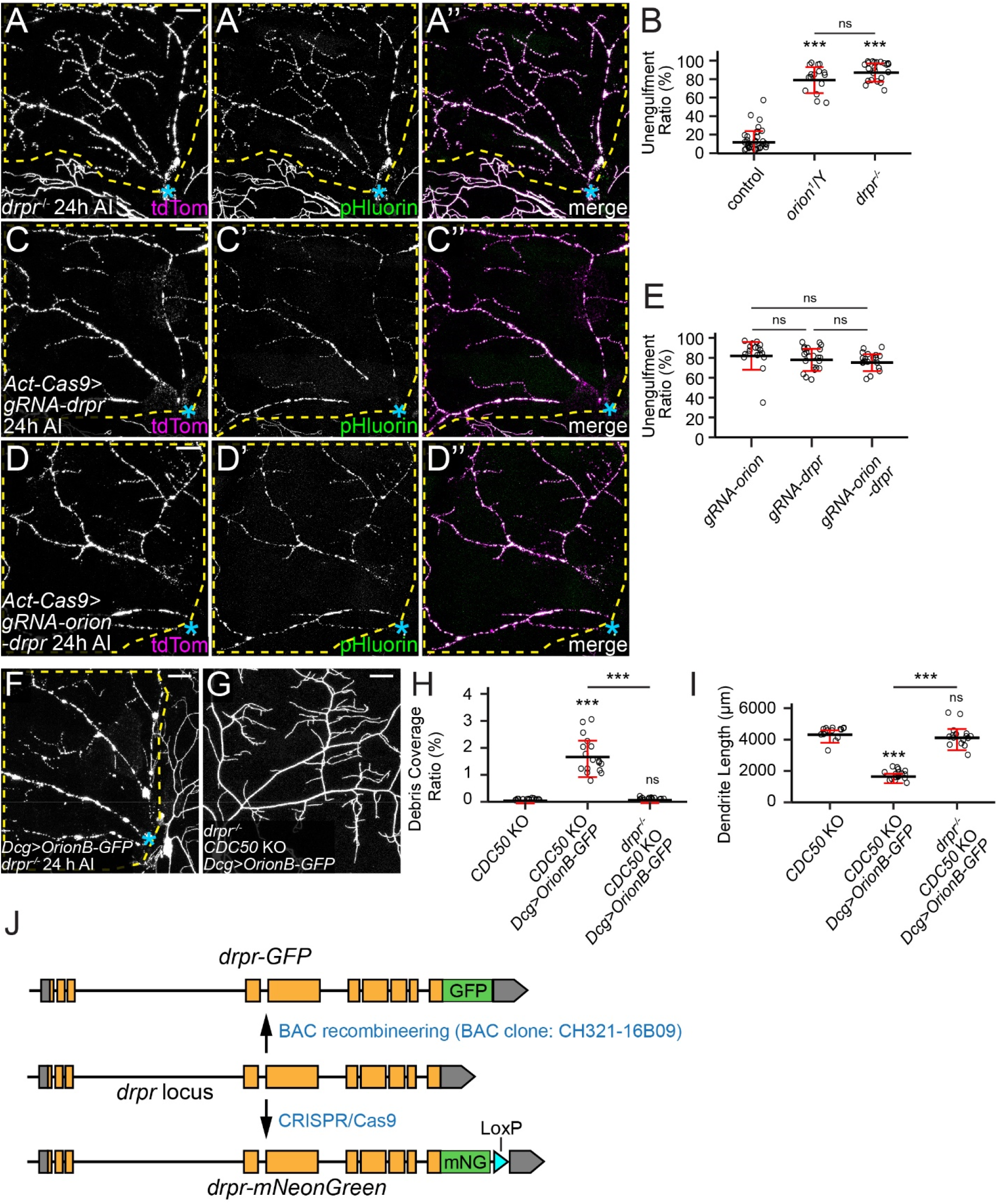
Orion recruits epidermal Drpr to injured dendrites. (A-A”) Partial dendritic fields of a ddaC neuron in *drpr*^-/-^ at 24 hrs AI. (B) Quantification of unengulfment ratio of injured dendrites. n = number of neurons and N = number of animals: control and *orion^1^*/Y (same dataset as in Figure 1D); *drpr*^-/-^ (n = 23, N = 10). One-way ANOVA and Tukey’s test. (C-D”) Partial dendritic fields of ddaC neurons at 24 hrs AI with whole-body *drpr* KO (C-C”), and *drpr+orion* double KO (D-D”). (E) Quantification of unengulfment ratio of injured dendrites. n = number of neurons and N = number of animals: *orion* KO (same dataset as in Figure 2L); *drpr* KO (n = 23, N = 12); *orion+drpr* double KO (n = 19, N = 10). One-way ANOVA and Tukey’s test. (F) Partial dendritic fields of an injured ddaC neuron with OrionB-GFP OE in fat body in the *drpr*^-/-^ background at 24 hrs AI. (G) Partial dendritic fields of a *CDC50* KO neuron with OrionB-GFP OE in fat body in the *drpr*^-/-^ background. (H-I) Quantification of debris coverage ratio (H) and dendrite length (I). n = number of neurons and N = number of animals: *CDC50* KO and *CDC50* KO + *Dcg*>*OrionB-GFP* (96 hrs AEL, same dataset as in Figure 3K); *drpr*^-/-^ + *CDC50* KO + *Dcg*>*OrionB-GFP* (120 hrs AEL, n = 16, N = 8). For (H), Kruskal-Wallis (One-way ANOVA on ranks) and Dunn’s test, p-values adjusted with the Benjamini-Hochberg method; for (I), one-way ANOVA and Tukey’s test; ***p≤0.001; n.s., not significant. The significance level above each genotype is for comparison with the control. Black bar, mean; red bars, SD. (J) Schematic representation of Drpr-GFP and Drpr-mNG. Yellow boxes indicate exons, grey boxes indicate untranslated regions. Neurons were labeled by *ppk-MApHS* (A-A”, C-D”), and *ppk-CD4-tdTom* (F-G). Scale bars, 25 μm.

**Figure S5:**
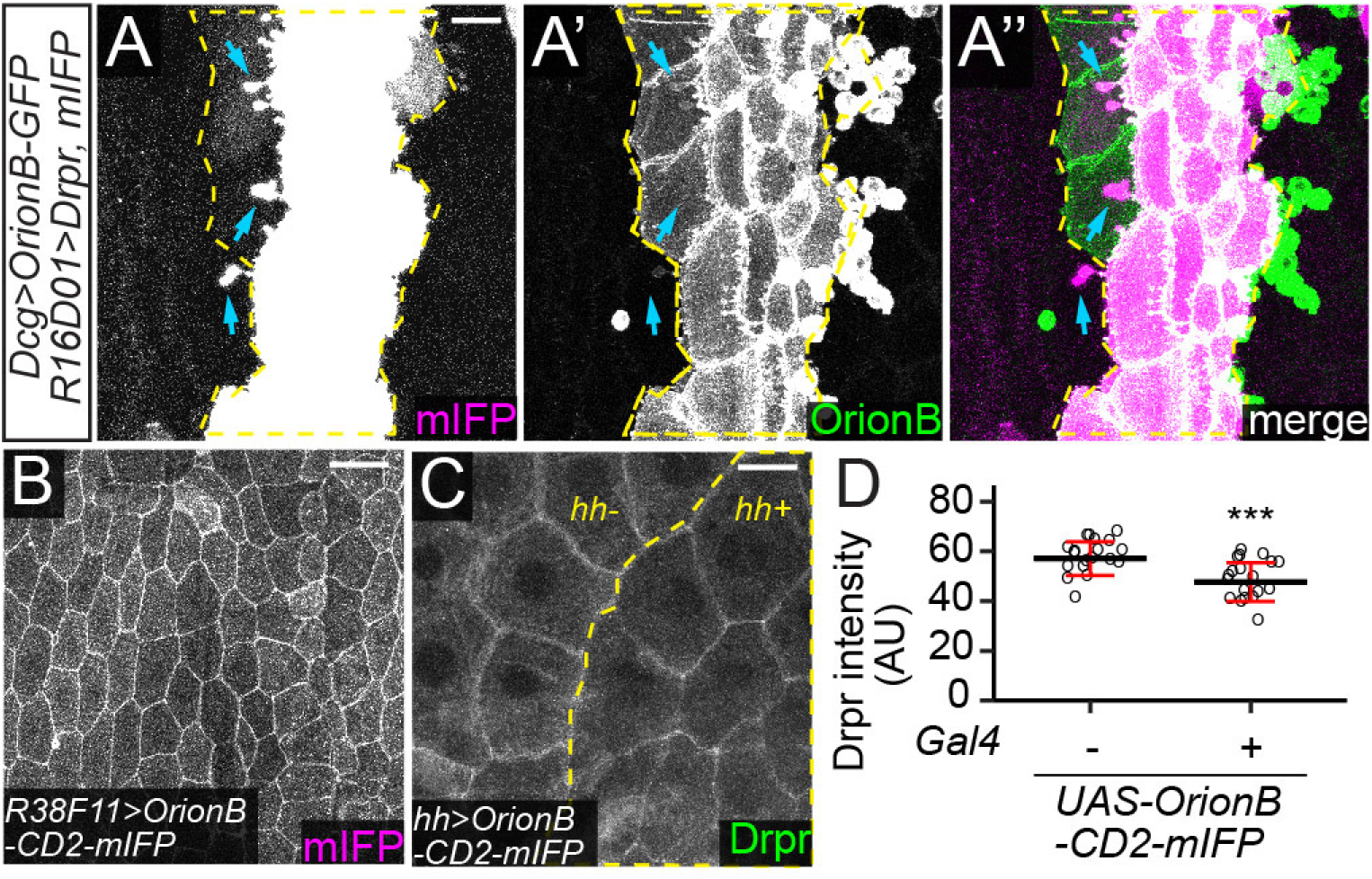
Orion mediates the interaction between PS and Drpr. (A-A”) Protrusions (blue arrows) extended from Drpr OE epidermal cells (labeled by mIFP) in the presence of fat body-derived OrionB-GFP. Yellow dash outlines: Drpr OE region. (B) OrionB-CD2-mIFP localization in epidermal cells driven by *R38F11-Gal4*. (C) Drpr localization in epidermal cells with OrionB-CD2-mIFP expressed in the *hh* domain. Yellow dash outlines: *hh-Gal4>OrionB-CD2-mIFP* region. (D) Quantification of stained Drpr levels in epidermal cells with or without OrionB-CD2-mIFP OE. n = number of measurements: w/o OrionB-CD2-mIFP OE (n = 19); w/ OrionB-CD2-mIFP OE (n = 19). Welch’s t-test; ***p≤0.001; black bar, mean; red bars, SD. Scale bars, 25 μm (A-A” and C), 50 μm (B).

**Figure S6:**
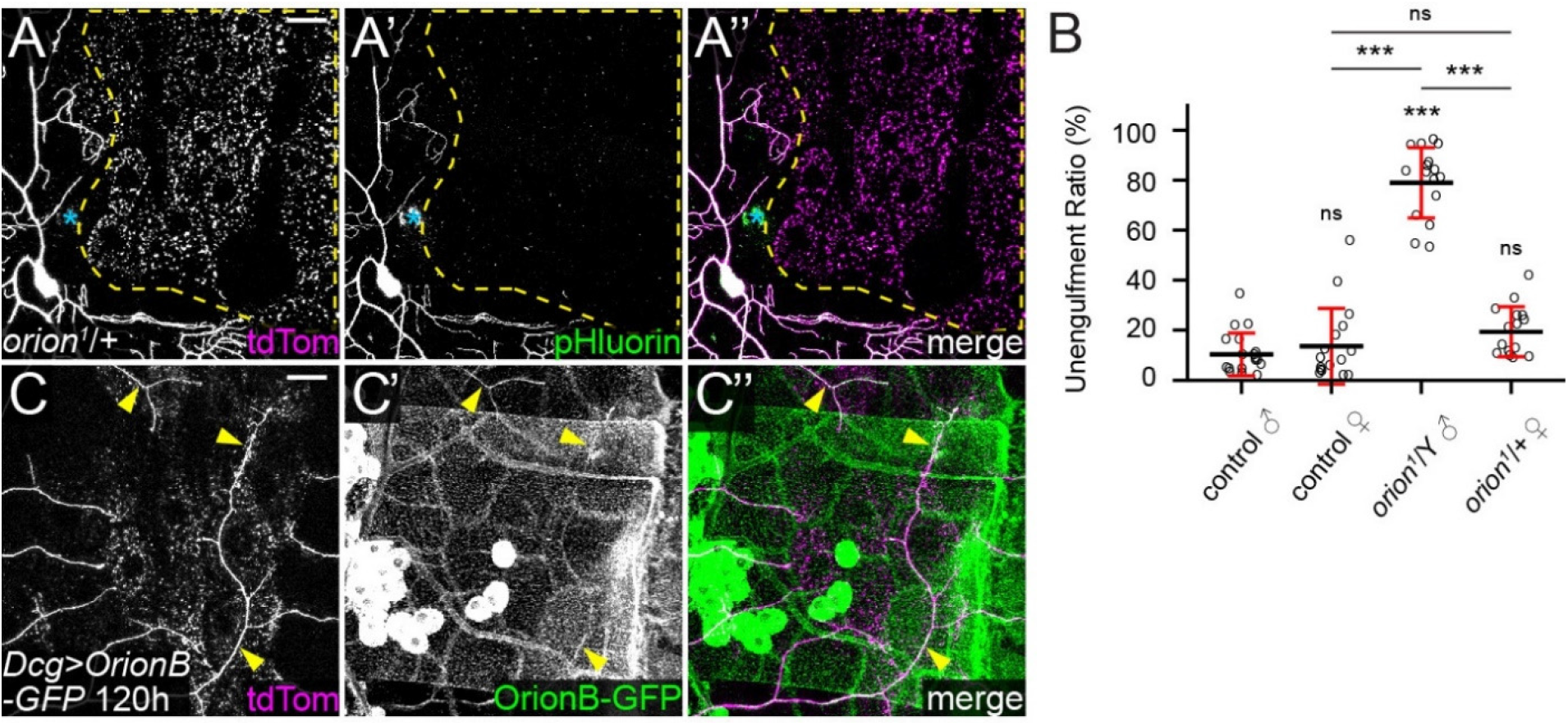
The Orion dosage determines the sensitivity of epidermal cells to PS-exposing dendrites. (A-A”) Partial dendritic fields of a ddaC neuron in an *orion^1^* heterozygous larva at 22 hrs AI. Yellow dash outlines: territories originally covered by injured dendrites; blue asterisks: injury sites. (B) Quantification of unengulfment ratio of injured dendrites. n = number of neurons and N = number of animals: control males, control females, and *orion^1^*/Y (same dataset as in Figure 1D); *orion^1^*/+ (n = 15, N = 8). One-way ANOVA and Tukey’s test; ***p≤0.001; n.s., not significant. The significance level above each genotype is for comparison with the control. Black bar, mean; red bars, SD. (C-C”) Partial dendritic fields of a wildtype neuron with OrionB-GFP OE in the fat body at 120 hrs AEL. Yellow arrowheads: OrionB-GFP colocalizing with dendrites. Neurons were labeled by *ppk-MApHS* (A-A”), and *ppk-CD4-tdTom* (C-C”). For all image panels, scale bars, 25 μm.

