## Supplemental Methods for "The *Drosophila* chemokine-like Orion bridges phosphatidylserine and Draper in phagocytosis of neurons"

### Supplementary Materials and Methods

#### Molecular cloning and transgenic flies

*orion-mNG2$_{11x4}$-V5-T2A-LexA*: Two copies of a gRNA spacer sequence targeting *orion* C-terminus and 3'UTR (Table S2) were cloned into pAC-CR7T-gRNA2.1-nlsBFP (Addgene 170515) according to published protocols (Koreman et al., 2021). The resulting plasmid was digested by PstI and NheI and assembled with four DNA fragments (through NEBuilder HiFi DNA assembly, New England Biolabs, Inc) to make an *orion* gRNA-donor vector. The four DNA fragments include a T2A-LexA-VP16 fragment that was PCR-amplified from pUC57-50-GS-FRTGFP2ALexAVP16-50 (Chen et al., 2014) (a gift from Larry Zipursky), an mNG2$_{11x4}$-V5 DNA fragment (synthesized by Integrated DNA Technologies, Inc.), and 5' and 3' homology arms (surrounding the stop codon of *orion*, ~1 kb each) that were PCR-amplified from the genomic DNA of *w$^{1118}$*.

*UAS-orionA-GFP:* OrionA coding sequence (CDS) was PCR-amplified from cDNA clone LD24308 (*Drosophila* Genomics Resource Center) and assembled with a superfolder GFP (sfGFP) fragment into pIHEU-MCS (Addgene 58375) (Sapar et al., 2018), resulting in pIHEU-orionA-GFP.

*UAS-orionB-GFP*: The first two exons of *orionB*, together with the first intron, was PCR-amplified from *w$^{1118}$* genomic DNA. The common CDS of OrionA and OrionB was PCR-amplified from LD24308. Both fragments were assembled into pIHEU-orionA-GFP to replace the orionA CDS, resulting in pIHEU-orionB-GFP.

*UAS-orionB-CD2-mIFP*: A pACU-CD2-mIFP plasmid was first constructed in pACU (Addgene 58373) (Han et al., 2011). The CDS of CD2-mIFP contains, from the N-terminus to the C-terminus, the rat CD2 CDS (AA24 to AA344), mIFP CDS (Yu et al., 2015), and Kir2.1 ER exit signal (Han et al., 2011). The OrionB CDS was then inserted before CD2 to make pACU-orionB-CD2-mIFP.

*UAS-orionB$^{AX3C}$-GFP*: An OrionB$^{AX3C}$ coding fragment was amplified from pENTR-orionB-AX$_3$C (Boulanger et al., 2021) and used to replace OrionB in pIHEU-orionB-GFP, resulting in pIHEU-orionB-AX$_3$C-GFP.

*UAS-orionB$^{AAY}$-GFP*: The RRY motif in OrionB coding sequence was changed into AAY by mutagenesis PCR. The mutated OrionB fragment was used to replace OrionB in pIHEU-orionB-GFP, resulting in pIHEU-orionB-AAY-GFP.

*UAS-orionB$^{1}$-GFP*: A G611D mutation was introduced into OrionB sequence by overlap-extension PCR. The mutated OrionB fragment was used to replace OrionB in pIHEU-orionB-GFP, resulting in pIHEU-orionB-G611D-GFP.

*LexAop-orionB-GFP*: The OrionB-sfGFP CDS was inserted into KpnI/XbaI sites of pAPLO vector (Poe et al., 2017).

*UAS-Drpr$^{\Delta Cyto}$*: The extracellular domain and transmembrane domain of Drpr (AA1 to AA827) was PCR-amplified from *UAS-drpr-I* (Logan et al., 2012) genomic DNA. An smFP-HA (non-fluorescent) fragment was PCR-amplified from pCAG-smFP-HA (Viswanathan et al., 2015) (a gift from Loren Looger). The two fragments were cloned into pACU through restriction cloning, resulting in pACU-DrprTM_smGFP(dark).

*UAS-smNG2$_{1-10}$*: An mNG2$_{1-10}$ fragment was synthesized (Integrated DNA Technologies, Inc.) and cloned into NheI/XbaI-digested pIHEU-sfGFPLactC1C2 (Sapar et al., 2018), resulting in pIHEU-smNG2(1-10).

*UAS-CDC50-T2A-ATP8A(E)*: The CDS of ATP8A isoform E (ATP8A(E)) was PCR-amplified from cDNA clone GH28327 (*Drosophila* Genomics Resource Center) and cloned into EcoRI/XbaI sites of pACU. The ATP8A(E) sequence is preceded by PacI and NheI sites and followed by a FLAG tag and a Kir2.1 ER exit signal (Sapar et al., 2018). In parallel, the CDC50 CDS was PCR-amplified from NB40 cDNA library (Brown and Kafatos, 1988) (a gift from Xinhua Lin) and cloned into EcoRI/XbaI sites of pACU. The CDC50 sequence is preceded by a PacI site and followed by BglII and NheI sites. A T2A fragment generated by annealed oligos was then inserted into the BglII/NheI sites. The CDC50-T2A (PacI/NheI) fragment was then released and cloned into PacI/NheI sites before ATP8A(E), resulting in pACU-CDC50-T2A-ATP8A(E).

*drpr-GFP*: The sfGFP CDS was inserted seamlessly before the stop codon of Drpr-PE in BAC clone CH321-16B09 according to published protocols of recombineering (Warming et al., 2005). Briefly, the *galK* CDS was first inserted before the stop codon of Drpr-PE in CH321-16B09 in bacterial strain SW102 through *galK*-mediated positive selection. The sfGFP CDS was then used to replace *galK* CDS in SW102 through *galK*-mediated negative selection. The resulting construct was transferred to bacterial strain EPI300 for propagation.

*drpr-mNG*: A *drpr-mNG* KI donor vector was constructed by assembling a pBluescript backbone and four DNA fragments. The four DNA fragments include the mNG CDS (Shaner et al., 2013), a 3xP3-GFP selection marker modified from pHD-DsRed (Addgene #51434), and 5' and 3' homology arms (surrounding the stop codon of Drpr-PE, ~1 kb each) that were PCR-amplified from CH321-16B09. A dual gRNA expression vector was constructed in pCFD4-U6.1_U6.3 (Port et al., 2014) to target the C-terminus and 3' UTR of *drpr* (Table S2).

*gRNA-orion* and *gRNA-orion-drpr*: A dual gRNA vector targeting *orion* and a quadruple gRNA vector targeting both *orion* and *drpr* were constructed in pAC-U63-QtgRNA2.1-BR (Addgene 170513) according to published protocols (Koreman et al., 2021).

Transgenic constructs were injected by Rainbow Transgenic Flies to transform flies through φC31 integrase-mediated integration into attP docker sites.

**Generation of KI flies**

To generate *drpr-mNG*, *drpr* KI donor vector and gRNA-expression vector were co-injected in *Act-Cas9* embryos. Adult flies from injected embryos were crossed to *w1118; TM3/TM6B*. The progeny was screened for green fluorescence in the adult eye. GFP-positive candidates were crossed to *$y^1$ $w^{67c23}$ $Cre(y+)^{1b}$; D/TM3, $Sb^1$* (BDSC, #851) to remove 3xP3-GFP. GFP-negative candidates were made isogenic and the mNG insertion was confirmed by genomic PCR and sequencing.

To generate *orion-mNG2$_{11x4}$-V5-T2A-LexA*, the *orion* gRNA-donor vector was injected into *$y^1$ nos-$Cas9^{ZH-2A}$ $w^*$* (BDSC, #54591) embryos. Adult flies from injected embryos were crossed to *$y^1$ $w^*$; 13XLexAop2-$6XGFP^{attP2}$/TM6B* (BDSC, #52266). GFP-positive female candidates from the progeny were collected to cross with *$y^1$ $w^*$; TM3/TM6B*. The adult progeny was then screened for GFP-positive males that were white-eyed and RFP-negative (i.e. having no *nos-$Cas9^{ZH-2A}$*). The males were then crossed to FM6 to remove *13XLexAop2-$6XGFP^{attP2}$* and to establish isogenic stocks. The -mNG211x4-V5-T2A-LexA insertion is verified by genomic PCR and sequencing.

**CRISPR-TRiM**

The efficiency of transgenic gRNA lines was validated by the Cas9-LEThAL assay (Poe et al., 2019). Homozygous males of each gRNA line were crossed to *Act-Cas9 w lig4* (BDSC, #58492) homozygous females. *gRNA-orion* crosses yielded viable female progeny and male lethality between 3$^{rd}$ instar larvae to prepupae; *gRNA-drpr* crosses resulted in lethality in late pupae; *gRNA-orion-drpr* crosses yielded viable female progeny and male lethality before wandering 3$^{rd}$ instar larvae. These results suggest that all gRNAs are efficient.

C4da-specific gene knockout was carried out using *ppk-Cas9* (Poe et al., 2019). Tissue-specific knockout in da neuron precursor cells was carried out with *SOP-Cas9* (Poe et al., 2019). Tissue-specific knockout in pan-epidermal cells was carried out using *shot-Cas9* (Ji et al., 2022). Tissue-specific knockout in epidermal cells in the posterior half of each segment was carried out using *hh-Cas9* (Poe et al., 2019). Whole-animal knockout was carried out using *Act-Cas9* (Port et al., 2014).

**Live imaging**

Animals were reared at 25°C in density-controlled vials (60-100 embryos/vial) on standard yeast-glucose medium (doi:10.1101/pdb.rec10907). Larvae at 96 hours AEL (3$^{rd}$ instar larval stage) or stages specified were mounted in 100% glycerol under coverslips with vacuum grease spacers and imaged using a Leica SP8 microscope equipped with a 40X NA1.30 oil objective. Larvae were lightly anesthetized with isoflurane before mounting. For consistency, we imaged dorsal ddaC neurons from A1-A3 segments (2-3 neurons per animal) on one side of the larvae. Unless stated otherwise, confocal images shown in all figures are maximum intensity projections of z stacks encompassing the epidermal layer and the sensory neurons beneath, which are typically 8–10 μm for 3$^{rd}$ instar larvae.

*Injury assay*

Injury assay at the larval stage was done as described previously (Sapar et al., 2018). Briefly, larvae at 84 hrs AEL were lightly anesthetized with isoflurane, mounted in a small amount of halocarbon oil under coverslips with grease spacers. The laser ablation was performed on a Zeiss LSM880 Confocal/Multiphoton Upright Microscope, using a 790 nm two-photon laser at primary dendrites of ddaC neurons in A1 and A3 segments. Animals were recovered on grape juice agar plates following lesion for appropriate times before imaging.

*Long-term time-lapse imaging*

Long-term time-lapse imaging at the larval stage was done as described previously (Ji et al., 2022; Sapar et al., 2018). Briefly, a layer of double-sided tape was placed on the coverslip to define the position of PDMS blocks. A small amount of UV glue was added to the groove of PDMS and to the coverslip. Anesthetized larvae were placed on top of the UV glue on the coverslip and then covered by PDMS blocks with the groove side contacting the larva. Glue was then cured by 365nm UV light. The coverslip with attached PDMS and larvae was mounted on an aluminum slide chamber that contained a piece of moistened Kimwipes (Kimtech Science) paper. Time-lapse imaging was performed on a Leica SP8 confocal equipped with a 40x NA1.3 oil objective and a resonant scanner at digital zoom 0.75 and a 3-min interval. For imaging after ablation, larvae were pre-mounted in the imaging chamber and subjected to laser injury. The larvae were then imaged 1-2 hours after ablation.

*Pinching assay*

Larvae at 96 hrs AEL were lightly anesthetized with isoflurane. Gentle pinching was performed at A2 or A3 segment and near the dorsal midline of larvae using a pair of forceps (DUMONT # 3, Fisher Scientifics) without cracking the cuticle. The pinched larvae were imaged after a 2-hr recovery.

**Immunohistochemistry**

Immunostaining of *Drosophila* larvae was performed as previously described (Poe et al., 2017). Briefly, $3^{rd}$ instar larvae were dissected in cold PBS, fixed in 4% formaldehyde/PBS for 20 min at room temperature (RT), and stained with the proper primary antibodies (Table S1) for 2 hrs at RT and subsequent secondary antibodies (Table S1) for 2 hrs at RT.

**Protein purification**

All the steps were carried out either on ice or at 4°C. 2 mL Ni-NTA Resin (ThermoFisher) was washed with lysis buffer (50 mM Tris-HCl (pH 7.5), 150 mM NaCl, 10% glycerol, 10 mM imidazole) three times for 5 min at 800x g. 125μL Ni-NTA Resin was washed with 1 mL PBS for three times at 800x g for 5 min. Purified 6xHis nanobody was thawed on ice and incubated with Ni-NTA Resin on a rotor for 1 hr. 50 *Drosophila* larvae (*Dcg-Gal4>UAS-OrionB-GFP*) were frozen at –80°C and thawed on ice. 300 μL of lysis buffer and 200 μL of 10 x protease inhibitor cocktail (Promega), 10 μL AEBSF (200x) (ThermoFisher) and 5 ceramic beads (Omni International) were added to the tube and homogenized using a beater (Omni International) for 3 times (30s beating, 30s on ice) at level 4. Another 500μL lysis buffer, 10 μL 10% TritonX-100 and 10 μL AEBSF (200x) (ThermoFisher) was added to the tube and centrifuged at 10,000x g for 10 min. 700 μL supernatant was added to 2 mL Ni-NTA Resin and incubated for 1 hr. The mixture was centrifuged at 800x g for 5 min. The flowthrough was then incubated in Ni-NTA-6xHis-nanbody for another hour and centrifuged at 800x g for 5 min. The pellet was wash with lysis buffer three times and then eluted with elution buffer (50 mM Tris-HCl (pH 7.5), 150 mM NaCl, 10% glycerol, 250 mM imidazole). The elution fraction was snap frozen with liquid N2 and stored at –80°C.

**Liposome preparation**

Dehydrated lipids (Avanti Polar Lipids) were dissolved in chloroform and combined in molar ratios described in Table S3. Lipids were dried under vacuum for 2 hr and then rehydrated in HK buffer (25 mM HEPES, pH 7.4, 125 mM KOAc) at 37°C for overnight. Lipids were extruded through 400 nm filters (Whatman) using a miniextruder (Avanti Polar Lipids). Lipids were extruded using 19 passes through the filter and stored at 4°C. Liposomes were used within 1 week of extrusion.

**Liposome sedimentation assay**

A 40-μL reaction in a 9.5 × 38 mm Polyallomer centrifuge tube (Beckman Coulter cat #357448) containing 250 μM liposomes and ~1 μg purified GFP tagged protein in HK buffer (with 2 mM CaCl2) were incubated at RT for 10 min. Mixtures was spun at 4°C at 55000 rpm for 30 min. Supernatant was removed via pipette immediately and 8 μL of 6x SDS sample buffer was added. Pellets were washed with HK buffer and resuspended with 48 μL of 1x SDS sample buffer (made by diluting 6x SDS sample buffer in HK buffer). All samples were heated at 55°C for 5 min and visualized by SDS-Page/Western Blot.

**Western blot**

The proteins were separated by 4-20% SDS-PAGE gel (Bio-Rad) and transferred onto nitrocellulose membranes (LiCOR). After blocking with 5% non-fat milk in TBST (Tris-buffered saline with 0.1% Tween 20) at RT for 1 hr, membranes were washed 3 times with TBST for 15 min and incubated with primary antibodies (anti-GFP Rabbit IgG Antibody Fraction, Alexa Fluor® 488 Conjugate, Life Technologies) overnight. ChemiDoc was used for detecting Alexa Fluor® 488.

**Image analysis and quantification**

Image processing and analyses were done in Fiji/ImageJ or ilastik. For injured dendrites and pruned dendrites marked by *ppk-MApHS,* the pHluorin-positive pixel area in a region of interest (ROI) (ApH), tdTom positive pixel area in the ROI (Atom) were measured and the unengulfment ratio was calculated based on following formula: 100·ApH/Atom. Methods for tracing and measuring C4da neuron dendrite length have been previously described (Poe et al., 2017). Briefly, the images were segmented by Auto Local Threshold and reduced to single pixel skeletons before measurement of skeleton length by pixel distance. The dendrite debris measurement has been described previously (Sapar et al., 2018). Briefly, a region of interest (ROI) was generated by including a quadrant of a neuron's territory. Dendrite debris within the ROI was converted to binary masks based on fixed thresholds. Different thresholds were used for *ppk-C4-tdTom* and *ppk-Gal4 UAS-CD-tdTom* as they have different brightness. The debris pixel area (Adeb), and ROI area (AROI) were measured, and dendrite coverage ratio was calculated based on following formula: 100·ADeb/AROI. For measuring debris dispersion of injured dendrites, dendrite debris was segmented by Auto Threshold (the "Default" method) in a rectangular ROI that was previously covered by injured dendrites. The ROI was divided into 15x15-pixel squares. The debris spread index was the area ratio of all squares containing dendrite debris in the ROI. For measuring Orion-GFP and AV-mCard on dendrites, tdTom signals on dendrites were used to generate dendrite masks for measurement of GFP or mCard mean intensities within the masks. For measuring Drpr-GFP recruitment, tdTom signals on dendrites were used to generate dendrite masks to measure total dendrite area (Atot) and GFP-positive area (AGFP). The Drpr recruitment index was calculated based on the formula: AGFP/Atot. For V5 staining, Drpr staining and Orion-GFP binding on epidermal cells, signals on cell boundaries of epidermal cells were measured. For Orion-GFP variant intensities in hemolymph, the signal in a single optical section was measured. For Orion-GFP variant intensities in fat body, maximum projected image was measured.

**Statistical Analysis**

R was used to conduct statistical analyses and generate graphs. (*p < 0.05, **p < 0.01, and ***p < 0.001). Statistical significance was set at $p < 0.05$. Data acquisition and quantification were performed non-blinded. Acquisition was performed in ImageJ. Statistical analyses were performed using R. We used the following R packages: car, stats, multcomp for statistical analysis and ggplot2 for generating graphs. For the statistical analysis we ran the following tests, ANOVA (followed by Tukey's HSD) when dependent variable was normally distributed and there was approximately equal variance across groups. When dependent variable was not normally distributed and variance was not equal across groups, we used Kruskal-Wallis (followed by Dunn's test, p-values adjusted with Benjamini-Hochberg method) to test the null hypothesis that assumes that the samples (groups) are from identical populations. We used Welch's t-test for comparison between two groups. To check whether the data fit a normal distribution, we generated qqPlots to analyze whether the residuals of the linear regression model are normally distributed. We used the Levene's test to check for equal variance within groups. The quantification of percentages of injured dendrites showing different timings of AV binding was compared using Fisher's exact test.

*Replication*

For all larval and adult imaging experiments, at least 3 biological replications were performed for each genotype and/or condition.

**Table S1. Key Resource Table**

| REAGENT or RESOURCE | SOURCE | IDENTIFIER | ADDITIONAL INFORMATION |
|---|---|---|---|
| **Experimental Models: Organisms/Strains** | | | |
| *orion*$^{1}$ | (Boulanger et al., 2021) | | |
| *orion*$^{\Delta C}$ | (Boulanger et al., 2021) | | |
| *orion*$^{\Delta A}$ | (Boulanger et al., 2021) | | |
| *orion*$^{\Delta B}$ | (Boulanger et al., 2021) | | |
| *orion-mNG2*$_{11x4}$*-V5-T2A-LexA (orion*$^{KI}$*)* | this study | | *orion-mNG(11x4)-F2A-LexA::VP16*$^{7A-1}$ |
| *Act5C-Cas9* | Bloomington Drosophila Stock Center | RRID:BDSC _54590 | *Act5C-Cas9.P* |
| *ppk-MApHS* | (Han et al., 2014) | | *ppk-MApHS*$^{1}$ |
| *UAS-TMEM16F* | (Sapar et al., 2018) | | *UAS-TMEM16F(D430G)*$^{VK00016}$ |
| *ppk-Gal4* | (Han et al., 2012) | | *ppk-Gal4*$^{VK00037}$ |
| *21-7-Gal4* | (Song et al., 2007) | | *GawB*$^{21-7}$ |
| *Dcg-Gal4* | (Suh et al., 2006) | | |
| *UAS-AnnexinV-mCard* | (Sapar et al., 2018) | | *UAS-AnnexinV-mCard*$^{VK00037}$ |
| *UAS-CD4-tdTom* | (Han et al., 2011) | | *UAS-CD4-tdTom*$^{7M1}$ |
| *UAS-OrionB-GFP* | this study | | *UAS-orion(B)-sfGFP*$^{VK00018}$ |
| *UAS-OrionB-Myc* | (Boulanger et al., 2021) | | *UAS-orion-B-myc* |
| *UAS-CDC50-T2A-ATP8A(E)* | this study | | *UAS-CDC50-T2A-ATP8A(E)*$^{VK00016}$ |

| | | | |
|---|---|---|---|
| *LexAop-OrionB-GFP* | this study | | *LexAop-orion(B)-sfGFP*$^{VK37}$ |
| *ppk-Cas9* | (Poe et al., 2019) | | *ppk-Cas9*$^{7D}$ |
| *UAS-Drpr* | Bloomington Drosophila Stock Center | RRID:BDSC_67035 | *UAS-drpr[I]* |
| *drpr-GFP* | this study | | *drpr-sfGFP*$^{VK00037}$ |
| *Dcg-LexA* | this study | | |
| *UAS-Drpr*$^{Cyto}$ | this study | | *UAS-drprTM-smGFP.HA*$^{VK00018}$ |
| *UAS-OrionB-CD2-mIFP* | this study | | *UAS-Orion(B)-CD2-mIFP*$^{VK00019}$ |
| *UAS-ATP8A* | (Ji et al., 2022) | | *UAS-ATP8Acore*$^{VK00016}$ |
| *UAS-OrionB*$^{AX3C}$*-GFP* | this study | | *UAS-orion(B.CX3Cmut)-sfGFP*$^{VK00018}$ |
| *UAS-OrionB*$^{AAY}$*-GFP* | this study | | *UAS-orion(B.RRYmut)-sfGFP*$^{VK00018}$ |
| *UAS-OrionB*$^{1}$*-GFP* | this study | | *UAS-orion(B.G611D)-sfGFP*$^{VK00018}$ |
| *UAS-AVmut-GFP* | (Sapar et al., 2018) | | *UAS-AnnexinV(mut)-GFP*$^{VK00018}$ |
| *ppk-Cas9* | (Poe et al., 2019) | | *ppk-Cas9*$^{7D}$ |
| *gRNA-CDC50* | (Sapar et al., 2018) | | *gRNA-CDC50*$^{attP2}$ |
| *gRNA-Nmnat* | (Ji et al., 2022) | | *gRNA-NmnatVK00027* |
| *LexAop-GFPnls* | Bloomington Drosophila Stock Center | RRID:BDSC _29955 | *lexAop-2xhrGFP.nls*$^{3a}$ |
| *gRNA-orion* | this study | | *gRNA-orion(BR)*$^{VK00027}$ |
| *shot-Cas9* | (Ji et al., 2022) | | *shot-Cas9*$^{1A}$ |
| *SOP-Cas9* | (Poe et al., 2019) | | *[sc-E1]x8-Cas9*$^{3A}$ |
| *R16D01-Gal4* | Bloomington Drosophila Stock Center | RRID: BDSC_48722 | *R16D01-Gal4*$^{attP2}$ |
| *UAS-smNG*$_{1-10}$ | this study | | *UAS-smNG2(1-10)*$^{VK00027}$ |
| *ppk-LexA* | (Poe et al., 2017) | | *ppk-LexA.GAD*$^{3}$ |
| *LexAop-Wld*$^{S}$ | (Ji et al., 2022) | | *LexAop-WldS*$^{VK00027}$ |
| *ppk-CD4-tdTom* | (Han et al., 2011) | | *ppk-spGFP11-CD4-tdTom*$^{2}$ |
| *R16A03-LexA* | (Sapar et al., 2018) | | *R16A03-LexAp65*$^{VK00027}$ |
| *UAS-mIFP-T2A-HO1* | (Poe et al., 2017) | RRID: BDSC_64181 | *UAS-mIFP-T2A-HO1*$^{VK00005}$ |
| *UAS-GFP-LactC1C2* | (Sapar et al., 2018) | | *UAS-GFP-LactC1C2*$^{VK00018}$ |
| *drpr*$^{-}$ | (Sapar et al., 2018) | | *drpr*$^{indel3}$ |

| | | | |
|---|---|---|---|
| *gRNA-drpr* | (Ji et al., 2022) | | *gRNA-drpr(BR)*$^{VK00027}$ |
| *gRNA-orion-drpr* | this study | | *gRNA-orion-drpr(BR)*$^{VK00027}$ |
| *hh-Cas9* | (Poe et al., 2019) | | *R28E04-Cas9*$^{6A}$ |
| *drpr-mNG* | this study | | *drpr-mNeonGreen*$^{1008\text{-}15}$ |
| *UAS-Drpr* | (Logan et al., 2012) | | *UAS-drpr[I]:HA* |
| *R38F11-Gal4* | Bloomington Drosophila Stock Center | RRID: BDSC_50014 | *R38F11-Gal4*$^{attP2}$ |
| *gRNA-ATP8A* | (Sapar et al., 2018) | | *gRNA-ATP8A*$^{VK00019}$ |
| *hh-Gal4* | (Han et al., 2004) | | |
| *Dp(1:3)DC496* | Bloomington Drosophila Stock Center | RRID:BDSC_33489 | *PBac{DC496}*$^{VK00033}$ |
| *ppk-Gal4* | (Han et al., 2012) | | *ppk-Gal4*$^{1a}$ |
| *UAS-CD4-tdTom* | (Han et al., 2011) | | *UAS-CD4-tdTom*$^{VK00033}$ |
| *y*$^{1}$ *w*$^{67c23}$ *Cre(y+)*$^{1b}$*; D/TM3, Sb*$^{1}$ | Bloomington Drosophila Stock Center | RRID:BDSC_851 | |
| *y*$^{1}$ *nos-Cas9*$^{ZH\text{-}2A}$ *w*$^{*}$ | Bloomington Drosophila Stock Center | RRID:BDSC_54591 | |
| *13XLexAop2-6XGFP* | Bloomington Drosophila Stock Center | RRID:BDSC_52266 | *13XLexAop2-6XGFP*$^{attP2}$ |
| *Act-Cas9 w lig4* | Bloomington Drosophila Stock Center | RRID:BDSC_58492 | *y*$^{1}$ *Act5C-Cas9(RFP-)*$^{ZH\text{-}2A}$ *w*$^{1118}$ *DNAlig4*$^{169}$ |
| **Recombinant DNA** | | | |
| pAC-CR7T-gRNA2.1-nlsBFP | (Koreman et al., 2021) | RRID:Addgene_170515 | |
| pUC57-50-GS-FRTGFP2ALexAVP16-50 | (Chen et al., 2014) | | |
| LD24308 | *Drosophila* Genomics Resource Center | | |
| pIHEU-MCS | (Sapar et al., 2018) | RRID:Addgene_58375 | |
| pACU | (Han et al., 2011) | RRID:Addgene_58373 | |
| pENTR-orionB-AX3C | (Boulanger et al., 2021) | | |
| pAPLO | (Poe et al., 2017) | RRID:Addgene_112805 | |
| pCAG-smFP-HA | (Viswanathan et al., 2015) | RRID:Addgene_59759 | |

| | | | |
|---|---|---|---|
| GH28327 | *Drosophila* Genomics Resource Center | | |
| NB40 cDNA library | (Brown and Kafatos, 1988) | | |
| CH321-16B09 | BACPAC Resources Center | | |
| pGalK | (Warming et al., 2005) | | |
| pHD-DsRed | Addgene | RRID:Addgene_51434 | |
| pCFD4-U6.1_U6.3 | (Port et al., 2014) | RRID:Addgene_49411 | |
| pAC-U63-QtgRNA2.1-BR | (Koreman et al., 2021) | RRID:Addgene_170513 | |
| **Bacterial Strains** | | | |
| SW102 | (Warming et al., 2005) | | |
| EPI300 | Lucigen Corporation | Cat. # C300C105 | |
| **Antibody** | | | |
| Rat-Elav-7E8A10 (1:20) | Developmental Studies Hybridoma Bank | AB_528218 | |
| 8D12 anti-Repo (1:20) | Developmental Studies Hybridoma Bank | AB_528448 | |
| V5 Tag Antibody (R960-25) (1:400) | Thermo Fisher Scientific | AB_2556564 | |
| Rabbit anti Drpr polyclonal (1:100) | (Freeman et al., 2003) | | |
| Cy™5 AffiniPure Donkey Anti-Rat IgG (H+L) (1:200) | Jackson ImmunoResearch Labs | AB_2340671 | 712-175-150 |
| Alexa Fluor® 647 AffiniPure Donkey Anti-Mouse IgG (H+L) (1:200) | Jackson ImmunoResearch Labs | RRID: AB_2340862 | 715-605-150 |
| Alexa Fluor® 488 AffiniPure Donkey Anti-Rabbit IgG (H+L) (1:200) | Jackson ImmunoResearch Labs | RRID: AB_2313584 | 711-545-152 |
| Alexa Fluor® 488 AffiniPure Donkey Anti-Mouse IgG (H+L) (1:400) | Jackson ImmunoResearch Labs | RRID: AB_2340846 | 715-545-150 |
| anti-GFP Rabbit IgG Antibody Fraction, Alexa Fluor® 488 Conjugate | Life Technologies | Catalog # A-11122 | |
| **Software and Algorithms** | | | |

| | | | |
|---|---|---|---|
| Fiji | https://fiji.sc/ | RRID: SCR_002285 | |
| R | https://www.r-project.org/ | RRID: SCR_001905 | |
| Adobe Photoshop | Adobe | RRID:SCR_014199 | |
| Adobe Illustrator | Adobe | RRID:SCR_010279 | |
| ilastik | (Berg et al., 2019) | | |
| UCSF Chimera | (Pettersen et al., 2004) | | |
| **Other** | | | |
| NEBuilder® HiFi DNA Assembly Master Mix | New England Biolabs Inc. | #E2621 | |
| HisPur™ Ni-NTA Resin | ThermoFisher Scientific | 88221 | |
| 6xHis nanobody | Addgene | #49172 | |
| Protease Inhibitor Cocktail 50X | Promega | G6521 | |
| AEBSF Protease Inhibitor | ThermoFisher Scientific | 78431 | |
| 2.8 mm Ceramic beads bulk, 325 g | Omni International | 19-646 | |
| Bead ruptor 4 | Omni International | 25-010 | |
| TritonX-100 | Amresco | M143 | Proteomics Grade |
| 1,2-dioleoyl-sn-glycero-3-phosphocholine (DOPC) | Avanti Polar Lipids, Inc. | Cat# 850375 | |
| 1,2-dioleoyl-sn-glycero-3-phospho-L-serine (DOPS) sodium salt | Avanti Polar Lipids, Inc. | Cat# 840035 | |
| 1,1'-Dioctadecyl-3,3,3',3'-Tetramethylindotricarbocyanine Iodide (DiR'; DilC18(7)) | ThermoFisher Scientific | Cat# D12731 | |
| Whatman® Nuclepore™ Track-Etched Membranes | Millipore Sigma | WHA110657 | |
| Extruder Set With Holder/Heating Block | Avanti Polar Lipids, Inc. | 610000-1EA | |
| 9.5 × 38 mm Polyallomer centrifuge tube | Beckman Coulter | Cat #357448 | |
| 4–20% Mini-PROTEAN® TGX™ Precast Protein Gels, 12-well, 20 µl | Bio-Rad | #4561095 | |
| Odyssey® Nitrocellulose Membranes | LI-COR | P/N 926-31092 | |
| ChemiDoc Imaging System | Bio-Rad | 17001401 | |

**Table S2. gRNA target sequences.**

| Gene | Target sequence 1 | Target sequence 2 |
|---|---|---|

| | | |
|---|---|---|
| *orion (for KO)* | GAAGGGCAACTACACCCAGG | CATGTTTCGTCGGATCACAG |
| *orion (for KI)* | GATTCTAAAGCGGAGAGAAG | |
| *drpr (for KO)* | CCATGCCGTAGAATCCAGGT | ACGGACAAGGATGCGCCCAG |
| *drpr (for KI)* | AGAAATTTCGGACTGGAACT | GCCGGAACAGTCACTTCACC |

**Table S3 Composition of liposomes**

| Lipid | Composition |
|---|---|
| PC | DOPC 99%, DiR dye 1%; |
| PC+PS | DOPC 79%, DOPS 20%, DiR dye 1% |
